# A new *Polerovirus* species in plants from the *Asparagaceae* family defined by its RNA-silencing repressor protein P0

**DOI:** 10.1101/2024.09.10.608920

**Authors:** Rob J. Dekker, Wim C. de Leeuw, Marina van Olst, Wim A. Ensink, Selina van Leeuwen, Timo M. Breit, Martijs J. Jonker

## Abstract

To contribute to the discovery of the global virome, we explored plants from an urban botanic garden in the Netherlands for new and variant plant viruses. Analyzing RNA from 25 plants from the *Asparagaceae* family with both RNA-seq, as well as smallRNA-seq revealed in six plants the presence of a variant Polerovirus from the *Solemoviridae* family that shows an overall RNA-sequence identity of 93% to the known Ornithogalum Virus 5 (OV-5). Amino acid sequence comparison of the complex set of proteins produced by the new virus revealed that all but Protein-0, showed high similarity (> 91%) with those of the OV-5 virus. The similarity between the new P0 protein and the OV-5 P0 protein, however, was for all variants less than 83%, well below the ICTV species protein-based demarcation criterium. Hence, we named the new virus Asparagaceae Polerovirus 1 (AspPolV-1).

## Introduction

The number of distinct virus species is estimated from 10^7^ to 10^9^, (Koonin *et al*., 2023). Hence, charting the global virome requires a global effort (Carrol *et al*., 2018). We try to contribute to this effort by analyzing plants from a specific habitat: the urban botanic garden. For this study, we focused on plants from *Asparagales* family to investigate the occurrence and sequence characteristics of known and unknown plant viruses. An example is the genus Polerovirus of the *Solemoviridae* family that encompasses at least 26 species infecting a wide range of plants (LaTourrette *et al*., 2021).

In this report we describe a new Polerovirus species that was discovered next to a new Betaflexivirus species (Dekker *et al*., 2024a) and a new Phenuivirus species (Dekker *et al*., 2024b) in an analysis of *Asparagales* plants from an urban botanic garden in the Netherlands. The discovery and characterization of this new Polerovirus species is a small contribution to the understanding of the global virome.

## Material and methods

### Samples

Samples of leaves from 25 *Asparagales* plants were collected from Hortus Botanicus, a botanic garden in Amsterdam, the Netherlands, on February 14, 2019. Details about the plant genera can be found in Supplemental Table ST1.

### RNA isolation

Small-RNA was isolated by grinding a flash-frozen ±1 cm^2^ leaf fragment to fine powder using mortar and pestle, dissolving the powder in QIAzol Lysis Reagent (Qiagen) and purifying the RNA using the miRNeasy Mini Kit (Qiagen). Separation of the total RNA in a small (<200 nt) and large (>200 nt) fraction, including DNase treatment of the large RNA isolates, was performed as described in the manufacturer’s instructions. The concentration of the RNA was determined using a NanoDrop ND-2000 (Thermo Fisher Scientific) and RNA quality was assessed using the 2200 TapeStation System with Agilent RNA ScreenTapes (Agilent Technologies).

### RNA-sequencing

Barcoded smallRNA-seq and RNA-seq libraries were generated using a Small RNA-seq Library Prep Kit (Lexogen) and a TruSeq Stranded Total RNA with Ribo-Zero Plant kit (Illumina), respectively. The size distribution of the libraries with indexed adapters was assessed using a 2200 TapeStation System with Agilent D1000 ScreenTapes (Agilent Technologies). The smallRNA-seq libraries from samples S01 to S12 and from samples S14-S26 were clustered and sequenced at 2×75 bp and 1×75 bp, on a NextSeq 550 System using a NextSeq 500/550 Mid Output Kit v2.5 or a NextSeq 500/550 High Output Kit v2.5 (75 cycles or 150 cycles; Illumina), respectively. RNA-seq libraries were clustered and sequenced at 2×150 bp on a NovaSeq 6000 System using the NovaSeq 6000 S4 Reagent Kit v1.5 (300 cycles; Illumina).

### Bioinformatics analyses

Sequencing reads were trimmed using trimmomatic v0.39 (Bolger *et al*., 2014) [parameters: LEADING:3; TRAILING:3; SLIDINGWINDOW:4:15; MINLEN:19]. Mapping of the trimmed reads to the NCBI virus database was performed using Bowtie2 v2.4.1 (Langmead *et al*., 2012). Contigs were assembled from smallRNA-seq data using all trimmed reads as input for SPAdes De Novo Assembler (Prjibelski *et al*., 2020) with parameter settings: only-assembler mode, coverage depth cutoff 10, and kmer length 17, 18 and 21. Assembly of contigs from RNA-seq data was performed with default settings.

## Results and discussion

25 leaf samples of *Asparagales* plants from different genera were obtained from an urban botanic garden in the Netherlands (Supplementary Table ST1). After RNA isolation, RNAseq, smallRNA seq (Dekker, *et al*., 2024a and 2024b; Supplemental Table ST1), and extensive bioinformatics analyses, we discovered in sample S14 a new virus RNA sequence that was quite similar to two Polerovirus sequences in GenBank: Ornithogalum Virus 5 (OV-5, MN204612.1) and Polerovirus sp. isolate M (PolV-M, KY769755.1). The RNA sequence identity between these two accessions is 99.8%. The RNA identity between the new Polerovirus sequence and the OV-5 and PolV-M sequences is 93% (Table1).

Mapping of both smallRNA-seq and RNA-seq sequencing reads to the new Polerovirus sequences revealed that this virus was present in six samples: S02, S03, S05, S09, S14, and S24 (Supplemental Table ST1). Samples S05 and S24 appear to contain two Polerovirus variants (Supplemental File SF1). The RNA sequences of the Poleroviruses found in this experiment show high similarity (≥ 92.8%) to OV-5 (Table 1). Poleroviruses express a complex set of proteins (Figure 1; Walker *et al*., 2022; Delfrosse *et al*., 2021). Except for protein P0, all proteins showed high similarity (> 91%) with those of the OV-5 and PolV-M viruses (Table 1 and Supplemental File SF2). In contrast, there were five Polerovirus variants in which the protein P0 is just 81% to 83% similar to OV-5 (Table 1 and Figure 2). This means that the newly discovered virus is a new species of the Polerovirus genus considering the ICTV species protein-based demarcation criterium used to demarcate species of the *Solemoviridae* genus Polerovirus: “*Differences in amino acid sequence identity of any gene product of greater than 10%*” (Walker *et al*., 2022). As these samples originate from plants of three genera from the Asparagaceae family, we named the new virus Asparagaceae Polerovirus 1 (AspPolV-1).

**Table 1.**
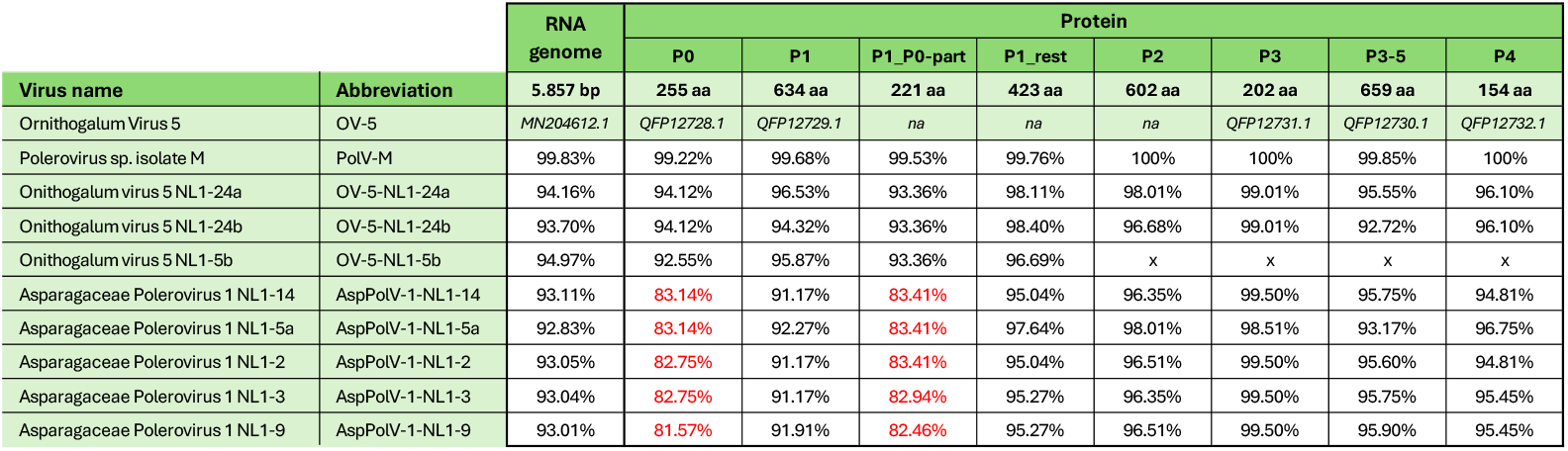
Genome and protein sequence comparison of Ornithogalum Virus 5 to related virus sequences. Sequence identities of the RNA genome and encoded proteins (P0-P4 and parts thereof) are shown for Polerovirus sp. isolate M, Ornithogalum virus 5 (OV-5) variants, and Asparagaceae Polerovirus 1 variants, compared to the reference sequence Ornithogalum Virus 5 (OV-5). The GenBank accession numbers for the OV-5 reference sequence are provided. Sequence identities for the P0 protein that are lower than 85% are highlighted in red.

**Figure 1.**
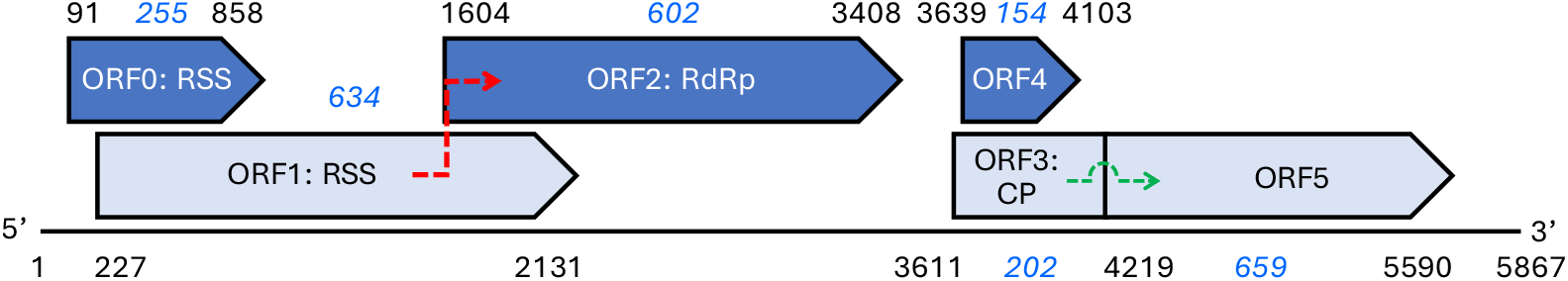
Schematic representation of the genome and proteins of *Asparagaceae Polerovirus 1* (AspPolV-1). Indicated are the positions, the translation start site and stop site (black), as well as the protein sizes (blue) of the proposed proteins (ORFs) of AspPolV-1. ORF2, coding for the RNA-dependent RNA polymerase (RdRp), is expressed as a fusion protein by a minus 1 ribosomal frameshift (red arrow). ORF5 is expressed as a fusion protein by a readthrough of the ORF3 stop codon at position 4,217 (green arrow). ORF0 codes for an RNA-silencing suppressor protein (RSS) and ORF3 codes for a coat protein (CP) (Mostert *et al*., 2020; Walker *et al*. 2022).

**Figure 2.**
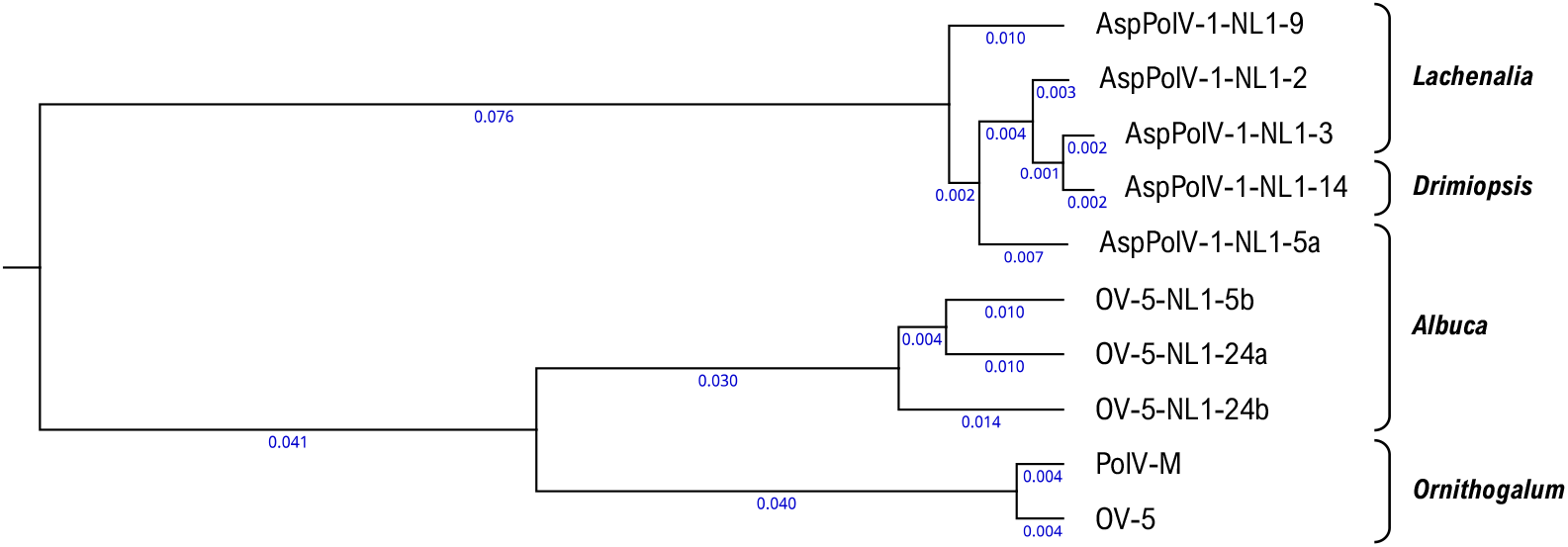
Phylogenetic guide tree constructed from the P0 protein sequences of AspPolV-1, OV-5, and their variants identified in this study. The tree was generated using the Clustal Omega 1.2.0 algorithm within the CLC Genomics Workbench 24.0.1 software. Branch lengths, indicating the evolutionary distance between sequences, are shown in blue below each branch. The genera of the host plants from which the viruses were derived are indicated in brackets next to the corresponding virus names.

It is of interest that the only protein showing distinct difference between these viruses is protein P0, the protein involved in suppression of the RNA resistance of host plants against virus infections (Baumberger *et al*., 2007, Csorba *et al*., 2010, Delfrosse *et al*., 2021). It is known that the amino acid sequence identities of Polerovirus P0 proteins are quite low, on average 37.06%, which also is reflected in varying silencing-suppressor activities between different virus isolates (Zhuo *et al*., 2014, LaTourrette *et al*., 2021, Delfrosse *et al*., 2021). The observed differences are definitively related to the P0 protein since the protein P1 differences mainly are present in the P1 part of the ORF that is overlapping with that of P0 (Table 1). Both OV-5 and PolV-M were isolated from symptomatic leaf material from plants of the genus Ornithogalum, whereas the AspPolV-1 virus was isolated from leaf material from plants of the genera Lachenalia, Albuca, and Drimiopsis (Figure 2; Supplemental Table ST1), which raises the possibility that this Polerovirus species variant adapted, via it P0 protein, to these genera, although one OV-5 variant (OV5-NL1-5b) was found in an Albuca plant. It would be interesting to investigate whether AspPolV-1 viruses are capable of infecting Ornithogalum plants.

## Acknowledgments

We would like to express our sincere gratitude to Sarina Veldman, Martin Smit, Iris van Kleinwee, and Reinout Havinga from the Hortus Botanicus in Amsterdam, The Netherlands, for their invaluable support in providing us with plant leaf material for this study. This research was directly and indirectly funded by the Swammerdam Institute for Life Sciences of the University of Amsterdam.

## Data availability

The raw sequence reads have been deposited in the NCBI Sequence Read Archive under BioProject accession number PRJNA1137160. The genome sequence of *Asparagaceae Polerovirus 1* isolate NL1-14 (AspPolV-1-NL1-14) was deposited in NCBI GenBank under accession PQ156406.

## Supplemental information

**Supplementary Table ST1.**
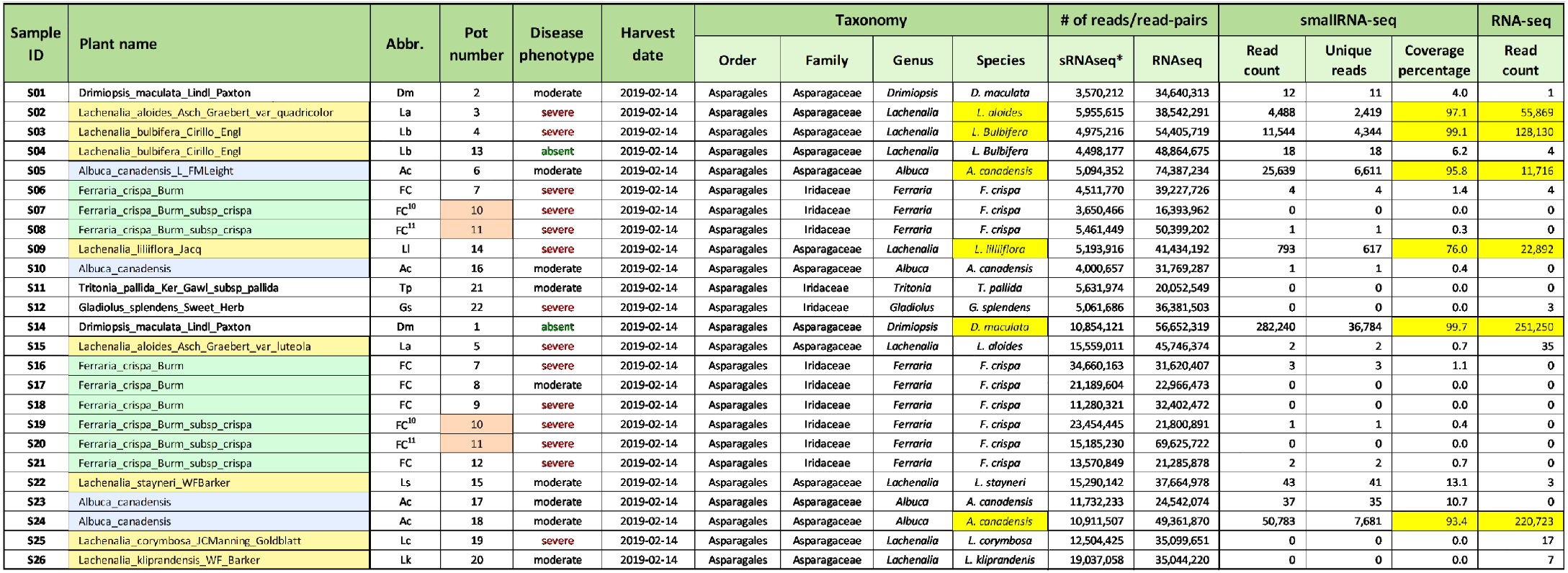
Overview of plant samples used for smallRNA and RNA-seq analyses. The table presents comprehensive details on each plant sample used in the smallRNA and RNA-seq analyses. Information includes the disease phenotype, harvest date, and taxonomic classification. For each sample, results are provided for the mapping of smallRNA-seq and RNA-seq reads to AspPolV-1. Specifically, the data includes the total number of reads, the count of unique reads, and the coverage percentage for smallRNA-seq, along with the number of read-pairs for RNA-seq. Samples from which the virus variants in this study were derived are highlighted in yellow. * The smallRNA-seq analyses were conducted in two separate experiments, which accounts for the observed variations in read counts.

### Supplemental File SF1. AspPolV virus RNA sequences

#### >AspPolV-1-NL1-2

TTCAGACGGTGTGCTCTTCCGATCCGAGAGAGCAGCTTTGAGCGCCTCCAACCCCCTTCCAAACCTTCAACTTAACCGTTTAAAAGTTGTATGTTTGTGATAACCTCTGAAGGATTCTTCTTGATAGATTCTTCTATCACTCCTTCTTTTGAGTTGTTGATTCTTCTTCTCCGCTCACTTGTTGCGTTCTGCATTTCTGCCCTACGCACCTCCTCGGTACCTTACGATGTCTTCTTCTTCAACTTTCATCGTTCTTTTCTTTTTGTTCTCCCTCTCCTCTGTGGTGTCCCCGTCCGCAACCTCTGCGACGGGCGATTTCGCGCCGCTCTCGACCTTCTCCCCTCTTTTCTGGAATGGGGCACTGTTACCCAATGCCACCCTCCTTTCCAGACTACCCGACGCCATGTCATTTTCGACATGCGCTTGTCCGGATCCGCCTCTCAATACATCACTCACTTACAACGAGTTGATGCGGATCTTCTGGGAAAAGGTCTCCTCCGACACCCAGAGCTTCTCCTCTCGCGCAATGGCAACCCTCAGCGAACACTCTGTTGCCTTGCTCGCCTCCGCTCGCAAGCACGCCTCCAATGCGCTCGAGTCCTTACTGTGGATTCCCGTCGCACTTTTTTGGACCAGCTTGTACTACGCGGGGGTAGTGGTGTGGGTTTGCTTGACCCAATATACTGTGATCGCGTTGGCGTATTTGTCGGTCGGTTGCTTCATAGCACTGGCTTGGAACGCCGTGATCTGGGTTTGTTCACGCTTTCCTATATTCCTTCTCTGTACACCCTTTTACGTCCTGAGGAGTGTTTGGAAGATTCTGTCATCCAAAAGCTTCTCCGTGAAGGTTGTCAATGAGAAAGCTGTGGACGGGTATGTGGATTATTCCATACCCCAGGACCCACCGAGAGGATCGACTGTACGCCTTCTTTATCCGAATGGCAACCCTTTGGGGTATGCTACTTGCGTACGGTTGTTCAATGGGGAGAACGCCCTCATAACCAGCTATCACTGCGTCGTTGAGGAAGGTGCTCTAGTTCACTCCACTCGCACCGGCAACAAATTGCCCCTTTCCCTGTTCAAACCCCTTTATGAGGATCAACGTGGTGATCTTTCGATCATGTGTGGTCCCCCTAATTGGGAAGGTTTGATGGGATGCAAGGGAGTTCATGCCGTGACTTGCGATCGCTTGGGACGCGGACCAGCCACTTTCTTCGTTTACGATAATAGCTGGAAAGCGAGGTCAGCCCAGATCGCTGGTCGCTATGAGGATTTCGCTCAGGTTCTTTCTAATACGGAACCTGGAGTGAGTGGGGCAGGGTATTTTAATGGCAAAACCCTGCTCGGTGTGCATAAGGGACACACCGACGTGGCTGAACACAACTTCAATTTAATGTCGCCACTCCCATGTCTCCCTGGTCTGACAGCCCCCATATACGTTTATGAGACCACAAACGTGCAGGGGAGAATCTTCGACGAGGACTTCGTTATACAATTTAAGAAGGGAAGCTGGTTCGATCAGATGATCGAACACGCTGAGAACCACGACGTGCTACCACCTTTGGTACGGCGTCCCGATGGTTCTCTCCGCTTTGCGGATGAGAAATCCCTCCCAATCGAGTTCTCGGGAAACGGAGCAGGCAGCGCCGTCTGCCCACCAAACGAGCCTGTCAAGACCCCGTCAGCCCCCCCCCCGATGCCCTCCCCCTCCCTTCTAGGAAGCTTTGTCGCTCCGTCCTCCGACACCCCTGTGACCGTCGGCGTTTCTTCGAACGCCTTGCAGGAAGGAGTCTTAAAGGCGTTGGTAGCGAAGTTCGACTTCCAGAAGGTAGAGCAGAATGTCGCAGAATTAATCGCTGCGAAAATGCTCTCCCGTCCTCAGCGAACGCGTGGTACCCGAGGGAGGAAGCGAAGCAATTCCAAAACTACTTCGCCTCCCTCTACAAGTGGTCAGAGTCAGCCCCCTCCGAAGCCCCAGGCTTCAAAACCGTCGGCAGGCTCCCCTCGTACTACCACCCCTCCCCAAAACGGGAAAGCAAATGGGGGAAAAAGCTCACCCGGCAATATCCCGAGCTGGGTGAGAAAACCACAGGCTTCGGCTGGCCGGGGGCTGGCGCAGAGGCGGAGCTGAAATCCCTTCAGCTCCAAGCGGCCAGGTGGCTCCAACGCGCGGAGACCGCCATCATTCCAAATCAGGCCGATCGGGAGCGCGTAATCGCTCGCACCGTTGACGCGTACTGTGCCGTTCATACGGAAGTACCATCGTGTGCGAGAACCAATGAGTTGACATGGCCAACGTTTCTTGAAGATTTTAAACGTGCCATTTGCTCGCTGGAGGTTGATGCCGGTATCGGCATCCCTTACATAGCCTACGGTCTCCCCACCCATCGTGGTTGGGTTGAGAATCCGTCTCTGCTACCAGTTCTAGCTCAACTCACATTTGCTCGTCTACAGAAGTTAGCGAATGTGAGCTTTGAAGCTTTGTCTGCAGAGGAGCTTGTAAGGTGTGGTCTTTGCGACCCTATCCGTCTTTTTATTAAGATGGAGCCTCATAAGCAGAGCAAACTCGATGAAGGTCGCTACCGCCTCATCATGAGTGTCTCATTGGTTGATCAATTGGTAGCCCGGGTTTTATTTCAAAATCAGAATCAGCGGGAGATCGCCTTGTGGAGGGCGATTCCAAGCAAACCCGGATTTGGACTATCCACTGACGAGCAAGTCGCGGACTTTCTTGCTTGTCTTGCTAAGGTAGTGGGAACTACACCTACTGAAGCTTGCAGCTGTTGGCGCGAGCTTATGATTCCCACAGACTGCTCCGGTTTCGACTGGAGCGTCGCCGCTTGGATGCTCGAGGATGAAATGGAGGTGAGAAATCGCCTGACCATCAATAATACCGAGCTGTGTCGCCGCCTTCGGGCCGGCTGGACGCATTGCATCGCTAATTCTGTGCTTTGCTTGTCCGACGGCACCTTACTCGCCCAGACGGTGCCTGGCGTGCAGAAGTCCGGGAGTTACAATACTTCTTCATCTAACTCCCGAATTCGCGTCATGGCAGCCTTTCACGCAGGGGCTTCCTGGGCGATGGCGATGGGCGATGATGCCCTCGAGTCCACAGACAGCCAAATCCCTGTCTATAAAAATCTTGGATTTAAAGTCGAGGTCGCGGAACAGTTGGAATTTTGTTCACATGTTTTTGTGCGTCCGGACCTCGCCATTCCGACCGGAACCAATAAGATGCTGTATAAGCTTCTATTTGGTTACGATCCGGAATGTGGGAACCTAGAGGTACTCTCCAATTACATTGCTGCATGTTGGAGTATCCTCAACGAGCTTCGATCTGATCCCGAACTCGTTTCCCGCCTCTACTCGTGGCTGATTCCAACGCAGTCACAAAAGAATACCGCGGAGTAGAGCAAATAGGAAAACACATAGCCTCACATACACAGTTGCAAGTGGAGGAGTGTTTAGTCTGTTACCACGCGACGTTAAACAATTGATTTTAAGTTTATTGCCGGTTTTGGGCTTGGCTTTTTAGCCGCCATACCGATTTCAGTCGTCGCGATTTATTTTATCTACCTAAAGATCTCAGCCAACGTACGCTCAATTGTTAATGAATACGGGCGGGGCTAGATCTAGAAATGGACGTCGTAGAGTTCGACTACCACGCCGCCTCCAGCGGCGTAGGCCAGCTCAGCCAATCGTTGTGGTCTCGGGACCCAACCAGATTCGACGCCGTCGACGACGAAGAGGAAATTCTTCACGTCGAGGAGCAAATGCAGCTCCAGGAAGAGGAGGCTCTCGTGAAACATTTGTTTTCACGAAGGATGACCTCGCGGGCAACACCTCTGGAAGTCTCACCTTCGGGCCGAGTCTTTCAGAGTGTCCAGCATTCGCGACAGGAATTCTCAAAGCCTACCATGAGTATCGTATCTCACAGTGTACTTTGGAGTTCATCTCCGAAGCCCCTTCCACGGCGTCCGGTTCAATCGCTTATGAGCTGGATGCACACTGCAAAATCTCCTCTCTCTCCTCAAAAATCAACAAGTTTGGAATCACTAAAGGAGGGAAGAGGAGCTTTGCGGCGTCCAAAATCAACGGGATAGAGTGGCATGATTCTTCTGAGGATCAGTTTCGCCTTTTATACAAAGGCAATGGCGCTTCCAATATCGCTGGCTCTTTTCGAGTCACCATTGTCGTCGCCACTCAAAACCCGAAATAGGTAGACGGCGAACCAGGGCCCAGTCCAGGGCCAGACCCCCCCCCTCCAGCACCAGCTCCCCAACCACACAAGCACGAACGTTTCATTGCTTATGTTGGTATCCCTATGCTTACAATTCAAGCGAAGGAGAATGATGACGGGATCATTCTACGATCACTCGGACCACAACGCATGAAGTATATTGAGGATGAAAATCAAAATTATACAAATATTGATTCTCAATATTACTCTCAAACCAGTGTCTCCGCTGTACCCATGTACTACTTCAATGTCCCTTCGGGCAAGTGGAGTGTTGATATCAGTTGTGAAGGTTATCAGTGCACTTCAAGCACAACTGATCCCAACCGTGGACGAAGCGATGGTTTGATAGCGTATTCAAATAGTTTCGATGATTATTGGAATGTGGGAGAAGCTGATGGTGTTATAATCACCAAGTTGACCAATGACAACACGTACAGGATGGGTCACCCCGATCTCGAGGTGAATTCCTGTCATTTTCGTGACGGTCAACTGTTGGAGCGTGACGCCACTATTTCCTTTCACGTTGAGGCTTCCAGTGATGGAAGATTCTTCTTGATCGGTCCCGCCATTCAGAAAATGGAAAAGTATAATTATACGATTTCGTATGGTGAGTGGACCGATCGTGATATGGAATTGGGTCTAATCACAGTTATTCTTGATGAACATCTTGAGGATTCTGGTTCGCGATTTGGTAGGATCCAACGGAGGAAGACACGAGATGGACACGTGCATCTTACCTCCATAGCGGAAAGACAACCGCTTACACAAACTCCTCGTTCTTTGCGAGTTCTCGATACTGGTTCGAGACATAAGTCCCTCTCACCTGAGAGGGCACGATCCCCCTCCCCCCTTAGGGAACCTTCACCACCTCCTCCGGAAAATCCGGAGATGTGGCAAGTGGATTGGCATGCTCCGAAATCTGGAGAACCTTTTGCTCCCTTGCCCCAGGATTTCAAACGTCCTGAGAGTGCGTTGAAGGGTGTCTTGCGTGAATCTCGCGGTAACACCGGAAATTATGAGCGAGATTACGCGGATAAGGAAAATTGGGATCCTTCAACTCTGGAGGACCCCACTCGCTTTGGCACTAACTGGACTTCTTCCCAGATTGCTGAAGCGCAAGAGCAGACGGATGCTAAACGCAAAACTCGTTTCTCTCTTGTTCCTCGTTCTTTTAAGGGAGGTTCCTCCTTAGGTGGAGGGAGCCTTACGGGAGGTTCTCTGCGTAATACCCTTCGGGGTCGCTTAGAGAAGTTGACTGCTGAGCAGCTGATCCAATATCATAGGATCAAGCAGCAGCAGGGATCTACCGTCGCCCAGATGTTCCTTACGGATTGCCTGGGTGACGGTTAACTTGCTCGTTTATTAATGAGCCCACACTGCAAGTAGCTTAGGCGAGTGCGTGTGGTTCCAAGTTTGGTCTACAATAAAGACTGGTCTTTCGGAGACGACCATAAATACATCTCCCCCAGTAAGTGTTCTGGGTTATCAAAAACCCGGGCAGTTTGTGGTCTGCCTATCAAAACCAACAGAAGGGTAGCAACCTTCCCGCCCCACTCGTTGTCTGCAGCGAGTGAGGGTTCCATTAGCTGGAGAGCAAACTTAGAGGGTAATTCCCCTT

#### >AspPolV-1-NL1-3

TTCAGACGGTGTGCTCTTCCGATCCGAGAGAGCAGCTTTGAGCGCCTCCAACCCCCTTCCAAACCTTCAACTTAACCGTTTAAAAGTTGTATGTTTGTGATAACCTCTGAAGGATTCTTCTTGATAGACTCTTCTATCACTCCTTCTTTTGAGTTGTTGATTCTTCTTCTCCGCTCACTTGTTGCGTTCTGCATTTCTGCCCTACGCACCTCCTCGGTACCTTACGATGTCTTCTTCTTCAACTTTCATCGTTCTTTTCTTTTTGTTCTCCCTCTCCTCTGTGGTGTCCCCGTCCGCAACCTCTGCGACGGGCGATTTCGCGCCGCTCTCGACCTTCTCCCCTCTTTTCTGGAATGGGGCACTGTTACCCAATGCCACCCTCCTTTCCAGACTACCCGACGCCATGTCATTTTCGACATGCGCTTGTCCGGATCCGCCTCTCAATACATCACTCACTTACAACGAGTTGATGCGGATCTTCTGGGAAAAGGTCTCCTCCGACACCCAGAGCTTCTCCTCTCGCGCAATGGCAACCCTCAGCGAATACTCTGTTGCCTTGCTCGCCTCCGCTCGCAAGCACGCCTTCAATGCGCTCGAGTCCTTACTGTGGATTCCCGTCGCACTTTTTTGGACCAGCTTGTACTACGCGGGGGTAGTGGTGTGGGTTTGCTTGACCCAATATACTGTGATCGCGTTGGCGTATTTGTCGGTCGGTTGCTTCATAGCACTGGCTTGGAACGCCGTGATCTGGGTTTGTTCACGCTTTCCTATATTCCTTCTCTGTACACCCTTTTACGTCCTGAGGAGTGTTTGGAAGATTCTGTCATCCAAAAGCTTCTCCGTGAAGGTTGTCAATGAGAAAGCTGTGGACGG GTATGTGGATTATTCCATACCCCAGGACCCACCGAGAGGATCGACTGTACGCCTTCTTTATCCGAATGGCAACCCTTTGGGGTATGCTACTTGCGTACGGTTGTTCAATGGGGAGAACGCCCTCATAACCAGCTATCACTGCGTCGTTGAGGAAGGTGCTCTAGTTCACTCCACTCGCACCGGCAACAAATTGCCCCTTTCCCTGTTCAAACCCCTTTATGAGGATCAACGTGGTGATCTTTCGATCATGTGTGGTCCCCCTAATTGGGAAGGTTTGATGGGATGCAAGGGAGTTCATGCCGTGACTTGCGATCGCTTGGGACGCGGACCAGCCACTTTCTTCGTTTACGATAATAGCTGGAAAGCGAGGTCAGCCCAGATCGCTGGTCGCTATGAGGATTTCGCTCAGGTTCTTTCTAATACGGAACCTGGAGTGAGTGGGGCAGGGTATTTTAATGGTAAAACCCTGCTCGGTGTGCATAAGGGACACACCGACGTGGCTGAACACAACTTCAATTTAATGTCGCCACTCCCATGTCTCCCTGGTCTGACAGCCCCCATATACGTTTATGAGACCACAAACGTGCAGGGGAGAATCTTCGACGAGGACTTCGTTATACAATTTAAGAAGGGAAGCTGGTTCGATCAGATGATCGAACACGCTGAGAACCACGACGTGTTACCACCTTTGGTACGGCGTCCCGATGGTTCTCTCCGCTTTGCGGATGAGAAATCCCTCCCAATCGAGTTCTCGGGAAACGGAGCAGGCAGCGCCGTCTGCCCACCAAACGAGCCTGTCAAGACCCCGTCAGCCCCCCCCCCGATGCCCTCCCCCTCCCTTCTAGGAAGCTTTGTCGCTCCGTCCTCCGACACCCCTGTGACCGTCGGCGTTTCTTCGAACGCCTTGCAGGAAGGAGTCTTAAAGGCGTTGGTAGCGAAGTTCGACTTCCAGAAGGTAGAGCAGAATGTCGCAGAATTAATCGCTGCGAAAATGCTCTCCCGTCCTCAGCGAACGCGTGGTACCCGAGGGAGGAAGCGAAGCAATTCCAAAACTACTTCGCCTCCCTCTACAAGTGGTCAGAGTCAGCCCCCTCCGAAGCCCCAGGCTTCAAAACCGTCGGCAGGCTCCCCTCGTACTACCACCCCTCCCCAAAACGGGAAAGCAAATGGGGGAAAAAGCTCACCCGGCAATATCCCGAGCTGGGTGAGAAAACCACAGGCTTCGGCTGGCCGGGGGCTGGCGCAGAAGCGGAGCTGAAATCCCTTCAGCTCCAAGCGGCCAGGTGGCTCCAACGCGCGGAGACCGCCATCATTCCAAATCAGGCCGATCGGGAGCGCGTAATCGCTCGCACCGTTGACGCGTACTGTGCCGTTCATACGGAAGTACCATCGTGTGCGAGAACCAATGAGTTGACATGGCCAACGTTTCTTGAAGATTTTAAACGTGCCATTTGCTCGCTGGAGGTTGATGCCGGTATCGGCATCCCTTACATAGCCTACGGTCTCCCCACCCATCGTGGTTGGGTTGAGAATCCGTCTCTGCTACCAGTTCTAGCTCAACTCACATTTGCTCGTCTACAGAAGTTAGCGAATGTGAGTTTTGAAGCTTTGTCTGCAGAGGAGCTTGTAAGGTGTGGTCTTTGCGACCCTATTCGTCTTTTTATTAAGATGGAGCCTCATAAGCAGAGCAAACTCGATGAAGGTCGCTACCGCCTCATCATGAGTGTCTCATTGGTTGATCAATTGGTAGCCCGGGTTTTATTTCAAAATCAGAATCAGCGGGAGATCGCCTTGTGGAGGGTGATTCCAAGCAAACCCGGATTTGGACTATCCACTGACGAGCAAGTCGCGGACTTTCTTGCTTGTCTTGCTAAGGTAGTGGGAACTACACCTACTGAAGCTTGCAGCTGTTGGCGCGAGCTTATGATTCCCACAGACTGCTCCGGTTTCGACTGGAGCGTCGCCGCTTGGATGCTCGAGGATGAAATGGAGGTGAGAAATCGCCTGACCATCAATAATACCGAGCTGTGTCGCCGCCTTCGGGCCGGCTGGACGCATTGCATCGCTAATTCTGTGCTTTGCTTGTCCGACGGCACCTTACTCGCCCAGACGGTGCCTGGCGTGCAGAAGTCCGGGAGTTACAATACTTCTTCATCTAACTCCCGAATTCGCGTCATGGCAGCCTTTCACGCAGGGGCTTCCTGGGCGATGGCGATGGGCGATGATGCCCTCGAGTCCACAGACAGCCAAATCCCTGTCTATAAAAATCTTGGATTTAAAGTCGAGGTCGCGGAACAGTTGGAATTTTGTTCACATGTTTTTGTGCGTCCGGACCTCGCCATTCCGACCGGAACCAATAAGATGCTGTATAAGCTTCTATTTGGTTACGATCCGGAATGTGGGAACCTAGAGGTACTCTCCAATTACATTGCTGCATGTTGGAGTATCCTCAACGAGCTTCGATCTGATCCCGAACTCGTTTCCCGCCTCTACTCGTGGCTGATTCCAACGCAGTCACAAAAGAATACCGCGGAGTAGAGCAAATAGGAAAACATATAGCCTCACATACACAGTTGCAAGTGGAGGAGTGTTTAGTCTGTTACCACGCGACGTTAAACAATTGATTTTAAGTTTATTGCCGGTTTTGGGCTTGGCTTTTTAGCCGCCATACCGATTTCAGTCGTCGCGATTTATTTTATCTACCTAAAGATCTCAGCCAACGTACGCTCAATTGTTAATGAATACGGGCGGGGCTAGATCTAGAAATGGACGTCGTAGAGTTCGACTACCACGCCGCCTCCAGCGGCGTAGGCCAGCTCAGCCAATCGTTGTGGTCTCGGGACCCAACCAGATTCGACGCCGTCGACGACGAAGAGGAAATTCTTCACGTCGAGGAGCAAATGCAGCTCCAGGAAGAGGAGGCTCTCGTGAAACATTTGTTTTCACGAAGGATGACCTCGCGGGCAACACCTCTGGAAGTCTCACCTTCGGGCCGAGTCTTTCAGAGTGTCCAGCATTCGCGACAGGAATTCTCAAAGCCTACCATGAGTATCGTATCTCACAGTGTACTTTGGAGTTCATCTCCGAAGCCCCTTCCACGGCGTCCGGTTCAATCGCTTATGAGCTGGATGCACACTGCAAAATCTCCTCTCTCTCCTCGAAAATCAACAAGTTTGGAATCACTAAAGGAGGGAAGAGGAGCTTTGCGGCGTCCAAAATCAACGGGATAGAGTGGCATGATTCTTCTGAGGATCAGTTTCGCCTTTTATACAAAGGCAATGGCGCTTCCAATATCGCTGGCTCTTTTCGAGTCACCATTGTCGTCGCCACTCAAAACCCGAAATAGGTAGACGGCGAACCAGGGCCCAGTCCAGGGCCAGACCCCCCCCCTCCAGCACCAGCTCCCCAACCACACAAGCACGAACGTTTCATTGCTTATGTTGGTATCCCTATGCTTACAATTCAAGCGAAGGAGAATGACGACGGGATCATTCTACGATCACTCGGACCACAACGCATGAAGTATATTGAGGATGAAAATCAAAATTATACAAATATTGATTCTCAATATTACTCTCAAACCAGTGTCTCCGCTGTACCCATGTATTACTTCAATGTCCCTTCGGGCAAGTGGAGTGTTGATATCAGTTGTGAAGGTTATCAGTGCACTTCAAGCACAACTGATCCCAACCGTGGACGAAGCGATGGTTTGATAGCGTATTCAAATAGTTCCGATGATTATTGGAATGTGGGAGAAGCTGATGGTGTTATAATCACCAAGTTGACCAATGACAACACGTACAGGATGGGTCACCCCGATCTCGAGGTGAATTCCTGTCATTTTCGTGACGGTCAACTGTTGGAGCGTGACGCCACTATTTCCTTTCACGTTGAGGCTTCCAGTGATGGAAGATTCTTCTTGATCGGTCCCGCCATTCAGAAAATGGAAAAGTATAATTATACGATTTCGTATGGCGAGTGGACCGATCGTGATATGGAATTGGGTCTAATCACAGTTATTCTTGATGAACATCTTGAGGATTCTGGTTCGCGATTTGGTAGGATCCAACGGAGGAAGACACGAGATGGACACGTGCATCTTACCTCCATAGCGGAAAGACAACCGCTTACACAAACTCCTCGTTCTTTGCGAGTTCTCGATACTGGTTCGAGACATAAGTCCCTCTCACCTGAGAGGGCACGATCCCCCTCCCCCCTTAGGGAACCTTCACCACCTTCTCCGGAAAATCCGGAGATGTGGCAAGTGGATTGGCATGCTCCGAAATCTGGAGAACCTTTTGCTCCCTTGCCCCAGGATTTCAAACGTCCTGAGAGTGCGTTGAAGGGTGTCTTGCGTGAAGCTCGCGGTAACACCGGAAATTATGAGCGAGATTACGCGGATAAGGAAAATTGGGATCCTTCAACTCTGGAGGACCCCACTCGCTTTGGCACTAACTGGACTTCTTCCCAGATTGCTGAAGCGCAAGAGCAGACGGATGCTAAACGCAAAACTCGTTTCTCTCTTGTTCCTCGTTCTTTTAAGGGAGGTTCCTCCTTAGGTGGAGGGAGCCTTACGGGAGGTTCTCTGCGTAATACCCTTCGGGGTCGCTTAGAGAAGTTGACTGCTGAGCAGCTGATCCAATATCATAGGATCAAGCAGCAGCAGGGATCTACCGTCGCCCAGATGTTCCTTACGGATTGCCTGGGTGACGGTTAACTTGCTCGTTTATTAATGAGCCCACACTGCAAGTAGCTTAGGCGAGTGCGTGTGGTTCCAAGTTTGGTCTACAATGAAGACTGGTCTTTCGGAGACGACCATAAATACATCTCCCCCAGTAAGTGTTCTGGGTTATCAAAAACCCGGGCAGTTTGTGGTCTGCCTATCAAAACCAACAGAAGGGTAGCAACCTTCCCGCCCCACTCGTTGTTTGCAGCGAGTGAGGGTTCCATTAGCTGGAGAGCAAACTTAGAGGGTAATTCCTCTTTGTTTGTTT

#### >AspPolV-1-NL1-5a

TTCAGACGGTGTGCTCTTCCGATCCGAGAGAGCAGCTTTGAGCGCCTCCAACTCCTTTCCAAACCTTCAACTTAACCGTCTAAAAGTTGTATGTTTGTGATTACCCCTGAAGGATTCTTCTTGATAGACTCTTCTATCACTCCTTCTTTTGAGTTGTTGATTCTTCTTCTCCGCTCACTTGTTGCGTTCTGCATTTCTGCCCTACGCACCTCCTCGGTACCTTACGATGTCTTCTTCTTCAACTTTCATCGTTCTTTTCTTTTTGTTCTCCCTCTCCTCTGTGGTGTCCCCGTCCGCAACCTCTGCGACGGGCGATTCCGCGCCGCTCTCGACCTTCTCACCGCTTTTCTGGAATGGGGCACTGTTACCCAATGCCACCCTCCTTTCCAGACTACCCGACGCCATGTCATTTTCGACATGCGCTTGTCCGGATCCGCCTCTCAATACATCACTCACTTACAACGAGTTGATGCGGATCTTCTGGGAAAAGGTCTCCTTCGACACCCAGAGCTTCTCCTCTCGCGCAACGGCAACCCTCAGCGAATACTCTGTTGCCTTGCTCGCCTCCGCTCGCAAGCACGCCTCCAATGCGCTCGAGTCCTTACTGTGGATTCCCGTCGCACTTTTTTGGACCAGCTTGTACTACGCGGGGGTAGTGGTGTGGGTTTGCTTGACCCAATATACTGTGATCGCGTTGGCGTATTTGTCGGTCGGTTGCTTCATAGCACTGGCTTGGAACGCCGCGATCTGGGTTTGTTCACGCTTTCCTATATTCCTTCTCTGTACACCCTTTTACGTCCTGAGGAGTGTTTGGAAGATTCTGTCATCCAAAAGCTTCTCCGTGAAGGTTGTCAATGAGAAAGCCGTGGACGGGTATGTGGATTATTCCATACCCCAGGACCCACCGAGAGGATCGACTGTACGCCTTCTTTATCCGAATGGCAACCCTTTGGGGTATGCTACTTGCGTACGGTTGTTCAATGGGGAGAACGCCCTCATAACCAGCTATCACTGCGTCGTTGAGGAAGGTGCTCTAGTTCACTCCACTCGCACCGGCAACAAATTGCCCCTTTCCCTGTTCAAACCCCTTTATGAGGATCAACGTGGTGATCTTTCGATCATGTGTGGTCCCCCTAATTGGGAAGGTTTGATGGGATGCAAGGGAGTTCATGCCGTGACTTGCGATCGCTTGGGACGCGGACCAGCCACTTTCTTCGTTTACGATAATAGCTGGAAAGCGAGGTCAGCTCAGATCGCTGGTCGCTATGAGGATTTCGCTCAGGTTCTTTCTAATACGGAACCTGGAGTGAGTGGGGCAGGGTATTTTAATGGCAAAACCCTGCTCGGTGTGCATAAGGGACACACCGACGTGGCTGAACACAACTTCAATTTAATGTCGCCACTCCCATGTCTCCCTGGTCTGACAGCCCCCATATACGTTTATGAGACCACAAACGTGCAGGGGAGAATCTTCGACGAGGACTTCGTTATACAATTTAAGAAGGGAAGCTGGTTCGATCAGATGATCGAACACGCTGAGAACCACGACGTGCTACCACCTTTGGTACGGCGTCCCGATGGTTCTCTCCGCTTTGCGGATGAGAAATCCCTCCCAATCGAGTTCTCGGGAAACGGAGCAGGCAGCGCCGTCTGCCCACCAAACGAGCCTGCCGAGACCCCGTCAGCCCCCCCCCCGATGCCCACCCCCTCCCTTCTAGGAAGCTTTGTCGCTCCACCCTCCGACACCCCTGCGACCGTCGGCGTTTCTTCAAACGCCTTGCAGGAAGGTGTCCTCAAGGCGTTGGTAGCGAAGTTCGACTTCCAGAAGGTCGAGCAGAATGTCGCAGAATTGATTGCTGCGAAAATGCTCTCCCGTCCTCAGCGAACGCGTGGTACCCGAGGGAGGAGACGAAGCAATTCCAAAACTACTTCGCCTCCCTCTACAAGTGGTCAGAGCCAGCCCCCTCTGAAGCCCCAGGCTTCAAAACCGTTGGCAGGCTCCCCTCGTACTACCACCCCTCCCCAAAACGAGAGAGCAAGTGGGGGAAAAAGCTCACCCGGCAACATCCCGAGCTGGGTGAGAAAACCCAAGGCTTCGGCTGGCCAGGGGCCGGCGCAGAAGCGGAGTTGAAAT CCCTTCAGCTCCAAGCGGCCAGGTGGCTCCAACGCGCGGAGACCGCCATCATTCCAAATCAGGCCGATCGGGAGCGCGTAATCGCTCGCACCGTTGACGCGTACTGTGCCGTTCATACGGAAGTACCATCGTGTGCGAGAACCAATGAGTTGACATGGCCAACGTTTCTTGAAGATTTTAAACGTGCCATTTGCTCGCTGGAGGTTGATGCCGGTATCGGCATCCCTTACATAGCCTACGGTCTCCCCACCCATCGTGGTTGGGTTGAGAATCCGTCTCTGCTACCAGTTCTAGCTCAACTCACATTTGCTCGTCTACAGAAGTTAGCGAATGTGAGTTTTGAAGCTTTGTCTGCAGAGGAGCTTGTAAGGTGTGGTCTTTGCGACCCTATCCGTCTTTTTATTAAGATGGAGCCTCATAAGCAGAGCAAACTCGATGAAGGTCGCTACCGCCTCATCATGAGTGTCTCATTGGTTGATCAATTGGTAGCCCGGGTTTTATTTCAAAATCAGAATCAGCGGGAGATCGCCTTGTGGAGGGCGATTCCAAGCAAACCCGGATTTGGACTATCCACTGACGAGCAAGTCGCGGACTTTCTTGCTTGTCTTGCTAAGGTAGTGGGAACTACACCTACTGAAGCTTGCAGCCGTTGGCGCGAGCTTATGATTCCCACAGACTGCTCCGGTTTCGACTGGAGCGTCGCCGCTTGGATGCTCGAGGATGAAATGGAGGTGAGAAATCGCCTGACCATCAATAATACCGAGCTGTGTCGCCGCCTTCGGGCTGGCTGGACGCATTGCATCGCTAATTCTGTGCTTTGCTTGTCCGACGGCACCTTACTCGCCCAGACGGTGCCTGGCGTGCAGAAGTCCGGGAGTTACAATACTTCTTCATCTAACTCCCGAATTCGCGTCATGGCAGCCTTTCACGCAGGGGCTTCCTGGGCGATGGCGATGGGCGATGATGCCCTCGAGTCCACAGACAGCCAAATCCCTGTCTATAAAAATCTTGGATTTAAAGTCGAGGTCGCGGAACAGTTGGAATTCTGTTCACACATTTTTGTGCGTCCGGACCTCGCCATTCCGACCGGGACCAACAAAATGCTGTACAAGCTTTTGTTTGGTTACGATCCGGAATGTGGGAACCTAGAGGTACTCTCCAACTACATTGCTGCATGTTGGAGCATCCTCAACGAGCTTCGATCTGATCCCGAGCTCGTTTCCCGTCTCTACTCGTGGCTGATTCCAACGCAGTCACAAAAGAATACCGCGGAGTAGAGAGAATAGGAAAACATAGCCTCACATACACAGTTGCAAGTGGAGGAGTGTTTTAGACTGTTACCACGCGACGTTAAACAATCGATTTTAAGTTTATTGCCGGTTTTGGGCTTGGCTTTTTAGCCGCCATACCGATTTCAGTCGTCGCGATTTATTTTATCTACCTAAAGATCTCAGCCAACGTACGCTCAATTGTTAATGAATACGGGCGGGGCTAGATCTAAAAATGGACGTCGTAGAGTTCGATTACCACGCCGCCTCCAGCGGCGTAGGCCAGCTCAGCCAATCGTTGTGGTCTCAGGACCCAACCAGATTCGACGCCGTCGACGACGAAGAGGAAATTCTTCACGTCGAGGAGCAAATGCAGCTTCAGGAAGAGGAGGCTCTCGTGAAACATTCGTTTTCACGAAGGATGACCTCGCGGGCAACACCTCTGGAAGTCTCACCTTCGGGCCGAGTCTTTCAGAGTGTCCAGCATTCGCGACAGGAATTCTCAAAGCCTACCATGAGTATCGTATCTCACAGTGTACTTTGGAGTTCATCTCCGAAGCCCCTTCCACGGCGTCCGGTTCAATCGCTTATGAGCTGGATGCACATTGCAAAATCTCCTCTCTCTCCTCGAAAATCAACAAGTTTGGAATCACCAAAGGAGGGAAGAGGAGCTTTGCGGCGTCCAAAATCAACGGGATAGAGTGGCATGATTCTTCCGAGGATCAATTTCGCCTTTTGTATAAAGGCAATGGCGCTTCTAATATTGCTGGCTCTTTTCGAGTCACTATTGTCGTCGCCACTCAAAACCCGAAATAGGTAGACGGCGAACCAGGGCCCAGCCCAGGGCCAGACCCCCCCCCTCCAGCACCAGCTCCCCAGCCACGTAAACATGAACGTTTCATTGCTTACGTTGGCATCCCTATGCTTACAATTCAAGCAAAGAAGAATGACGATGGGATCATTCTAAGGTCACTCGGACCCCAACGCATGAAGTATATTGAGGATGAAAATCAAAATTACACTAATATTGACTCCCAATACTACTCTCAAACCAATGTGTCCGCTGTCCCCATGTACTACTTCAATGTCCCAGCGGGCAAGTGGAGTGTTGATGTCAGTTGTGAAGGTTATCAGTGCACTTCAAGCACGACTGACCCCAACCGTGGACGAAGCGATGGTTTGATTGCGTATTCTAACAACTCCGATGACTATTGGAATGTGGGTGAAGCTGATGGTGTGATAATCACCAAGTTGACCAATGACAACACGTACAGGATGGGTCACCCCGATCTCGAGGTGAATTCCTGTCATTTTCGTGATGGTCAGTTGCTGGAGCGTGATGCCACGATTTCTTTTCACGTTGAGGCTTCCAGTGATGGAAGATTCTTCTTGATTGGTCCCGCCATACAGAAAATGGAAAAGTATAATTATACGATTTCGTATGGTGAGTGGACCGACCGTGATATGGAATTGGGTCTAATCACAGTTATTCTTGACGAACATCTTGAAGATTCTGGTTCGCGATTTGGTAAGATCCAACGGAGGAAGACACGAGATGGGCACGTGCGCCTTACCTCCGTAGCGGAAAGACAACCGCTTGTACAAACTCCTCGCCCCTTGCGAGTTCTCGATACTGGTTCGAGACGTAAGTCCCTCTCCCCGGAGAGGGCAAACTCCCCCCCCCCCCTTAGAGAACCTTCACCACCTCCTCCGGAAGATCCAGAGTTGTGGCAGGTCGGTTGGTACACCCCAAAATCTGGAGAACCACTGGCCCCTTTGCCCCAGGACTTTAAACGTCCTGAGAGTGTGTTGAAGGGTATCCTGCGTGAGTCCCGTGGTAACACCGGTGACTATGAAAAGGACTACGCGGATAAGGAAAATTGGGATCCTTCAACTTTGGAAGACCCAACTCGCTTCGGAACGAACTGGACAAGCTCCCAGATCGCCGAAGCACAAGAGCAGACGGATGCTAAACGCAAAACTCGTTTCTCTCTTGTTCCTCGTTCCTTTAAGGGAGGTTCCTCCATTAGTGGGGGAAGCTTATCGGGGGGTTCTCTTCGAAGCACTCTACGGAGTCGCTTGGAGAAGCTGACTGCTGAGCAGCAAATCCAATATCATCGGATTAAGCAGCAGCAGGGAGCTACCACCGCTCAGATGTTCCTTTCGGATTGCCTGGGTGATGGTTAACTTGCTCGTTTAATAACGAGCCCACACTGCTAGTAGCCTAGGCGAGCGCGTGTGGTTCCAAGTTTGGTCAACAATAAAGACTGGTCTTTCGGAGACGACCTGAAATACATCTCCCCCAGTAAGTGTTCTGGGTTATCAAAACCGGGGCAGTTAGTGGTCTGCCGATCAAAACCAACAGAAGGGTAGCAACCTTCCCACCCCCTGACACGTATTGTGTTAGTTGAGGGTTCCGTTAGCTTAGGAGAGCAAACTTAGAGGGTAATTCCTCTTTGTTTGTTT

#### >AspPolV-2-NL1-5b

TTCAGACGGTGTGCTCTTCCGATCCGAGAGAGCAGCTTTGAGCGCCTCCAACTCCTTTCCAAACCTTCAACTTAACCGTCTAAAAGTTGTATGTTTGTGATTACCCCTGAAGGATTCTTCTTGATAGACTCTTCTATCACTCCTTCTTTTGAGTTGTTGATTCTTCTTCTCCGCTCACTTGTCGCGTTCTGCTTTTCTGCTCAACGCACCTCCTCGGTTCCCGACGATGTCTTCTTCTACAATTTCCGTCGTTCTTTTCTTTTTGTTCTCCCTCTTCTTTGTGGCGTCCCCGTCCGCTACTTCAGCGACGGGCGCTTCCGCGCCACTCTCAACCTTCTCCCCGCTTTTCTGGAATGGGGCATTGTTACCCAATGCCACCCTCCTTTCCAGATTACCAAACGACATGTCATCTTCGACATGCGTTTGTTCGGATCCGCCTCTCGTTATATCACTCACTTACAACGAGTTGATGCGGATCTTCTCGGAGAAGGTCTGCTCCGACACTCGGAGCTTCTCCTCTCGCGCAATGGCAACCCTCAGCGAACACTCTGTTGCCTTGCTTACCTCCGCGCACAAGCACGCTTCCAGAGCGCTCGAGTCCTTTCTGTGGATTCCTGTCGCGCTTTTCTGGACCAGCTTGTATTACGTGGGGGTAGTAGTGTGGGTCTGCTTGACCCAGTACACTGCAATCGCATTGGCGTACTTATTGGTCGGTTGCTTCATAGCACTGGTTTGGAACGCCGCGATTTGGGTTTGTTCACGCTTTCCTTTATTCCTTCTTTGTACACCCTTCTACGTCCTGAAGAGTGTTTGGAAGACTCTGTCATTCAAACGCTCTTCCGTGAAGGTTGTGAATGAGAAAGCAGTGGACGGGTATGTGGATTATTCCATACCCCAAGACCCACCGAGAGGATCAACCGTGCGCCTTTTATACCCGAATGGAAATCCACTGGGGTATGCTACTTGCGTTCGGTTGTTTAATGGGGAGAACGCCCTCATAACCAGCTATCACTGCGTTATTGAGGAAGGTGTTCTGGTTCACTCCACTCGCACCGGCAACAAATTACCCCTTTCCCTTTTCAAGCCCCTTTATGAGGACCAACGTGGTGATCTTTCGATTATGTGCGGTCCTCCTAACTGGGAAGGTTTGATGGGATGCAAGGGAGTGCATGCCGTGACTTGCGATCGTTTGGGTCGTGGACCAGCTACCTTCTTTGTTTATGACAATAGCTGGAAGGCGAGGTCAGCCCAGATCGCTGGTCGCTATGAGGATTTCGCTCAGGTTCTTTCTAACACCGAACCTGGAGTGAGTGGGGCAGGGTACTTTAATGGTAAAACCCTGCTCGGTGTGCATAAGGGACACACCGACGTGGCTGAACACAATTTCAACTTGATGTCGCCACTCCCATGTCTCCCTGGTCTGACAGCCCCCATATACGTTTATGAAACCACGAACGTACAGGGGAGAATCTTCGACGAGGACTTTGTGGTCCAATTCAAGAAGGGAAGCTGGTTCGATCAGATGATCGAACACGCTGAGAACCACGATGTGCTACCTCCTTTGGTGCGCCGTCCTGATGGCTCTCTCCGCTTTGCGGATGAAAAATCCCTTCCCGCCAAATTCTCGGGAAACGAAGCTGGCAGCGCCGTCTGCCCACCAAACGAGCCTGTCAAGACCCCGTCAGCCCCCCCCCCGATGCCCTCCCCCTCCCTTCTAGGAAGCTTTGTCGCTCCACCCTCCGACACCCCTGC

#### >AspPolV-1-NL1-9

TTCAGACGGTGTGCTCTTCCGATCCGAGAGAGCAGCTTTGAGCGCCTCCAACCCCCTTCCAAACCTTCAACTTAACCGTTTAAAAGTTGTATGTTTGTGATAACCTCTGAAGGATTCTTCTTGATAGACTCTTCTATCACTCCTTCTTTTGAGTTGTTGATTCTTCTTCTCCGCTCACTTGTTGCGTTCTGCATTTCTGCCCTACGCACCTCCTCGGTACCTTACGATGTCTTCTTCTTCAACTTTCATCGTTCTTTTCTTTTTGTTCTCCCTCTCCTCTGTGGTGTCCCCGTCCGCAACCTCTGCGACGGGCGATTTCGCGCCGCTCTCGACCTTCTCCCCTCTTTTCTGGAATGGGGCACTGTTACCCAATGCCACCCTCCTTTCCAGACTACCCGACGCCATGTCATTTTCGACATGCGCTTGTCCGGATCCGCCTCTCAATACATCACTCACTTACAACGAGTTGATGCGGATCTTTTGGGAAAAGGTCTCCTCCGACACCCAGAGCTTCTCCTCTCGCGCAATGGCAACCCTCAGCGAATACTCTGTTGCCTTGCTCGCCTCCGCTCGCAAGCACGCCTCCAATGCGCTCGAGTCCTTACTGTGGATTCCCGTCGCACTTTTTTGTACCAGCTTGTACTACGCGGGGGTAGTGGTGTGGGTTTGCTTGACCCAATATACTGTGATCGCGTTGGCGTATTTGTCGGCCGGTTGCTTCATAGCACTGGCTTGGAACGCCATGATCTGGGTTTGTTCACGCTTTCCTATGTTCCTTCTCTGTACACCCTTTTACGTCCTGAGGAGTGTTTGGAAGATTCTGTCATCCAAAAGCTTCTCCGTGAAGGTTGTCAATGAGAAAGCCGTGGACGGGTATGTGGATTATTCCATACCCCAGGACCCACCGAGAGGATCGACTGTACGCCTTCTTTATCCGAATGGCAACCCTTTGGGGTATGCTACTTGCGTACGGTTGTTCAATGGGGAGAACGCCCTCATAACCAGCTATCACTGCGTCGTTGAGGAAGGTGCTCTAGTTCACTCCACTCGCACCGGCAACAAATTGCCCCTTTCCCTGTTCAAACCCCTTTATGAGGATCAACGTGGTGATCTTTCGATCATGTGTGGTCCCCCTAATTGGGAAGGTTTGATGGGATGCAAGGG AGTTCATGCCGTGACTTGCGATCGCTTGGGACGCGGACCAGCCACTTTCTTCGTTTACGATAATAGCTGGAAAGCGAGGTCAGCCCAGATCGCTGGTCGCTATGAGGATTTCGCTCAGGTTCTTTCTAATACGGAACCTGGAGTGAGTGGGGCAGGGTATTTTAATGGCAAAACCCTGCTCGGTGTGCATAAGGGACACACCGACGTGGCTGAACACAACTTCAATTTAATGTCGCCACTCCCATGTCTCCCTGGTCTGACAGCCCCCATATACGTTTATGAGACCACAAACGTGCAGGGGAGAATCTTCGACGAGGACTTCGTTATACAATTTAAGAAGGGAAGCTGGTTCGATCAGATGATCGAACACGCTGAGAACCACGACGTGCTACCACCTTTGGTACGGCGTCCCGATGGTTCTCTCCGCTTTGCGGATGAGAAATCCCTCCCAATCGAGTTCTCGGGAAACGGAGCAGGCAGCGCCGTCTGCCCACCAAACGAGCCTGTCAAGACCCCGTCAGCCCCCCCCCCGATGCCCTCCCCCTCCCTTCTAGGAAGCTTTGTCGCTCCGTCCTCCGACACCCCTGTGACCGTCGGCGTTTCTTCGAACGCCTTGCAGGAAGGAGTCTTAAAGGCGTTGGTAGCGAAGTTCGACTTCCAGAAGGTAGAGCAGAATGTCGCAGAATTAATCGCTGCGAAAATGCTCTCCCGTCCTCAGCGAACGCGTGGTACCCGAGGGAGGAAGCGAAGCAATTCCAAAACTACTTCGCCTCCCTCTACAAGTGGTCAGAGTCAGCCCCCTCCGAAGCCCCAGGCTTCAAAACCGTCGGCAGGCTCCCCTCGTACTACCACCCCTCCCCAAAACGGGAAAGCAAATGGGGGAAAAAGCTCACCCGGCAATATCCCGAGCTGGGTGAGAAAACCACAGGCTTCGGCTGGCCGGGGGCTGGCGCAGAAGCGGAGCTGAAATCCCTTCAGCTCCAAGCGGCCAGGTGGCTCCAACGCGCGGAGACCGCCATCATTCCAAATCAGGCCGATCGGGAGCGCGTAATCGCTCGCACCGTTGACGCGTACTGTGCCGTTCATACGGAAGTACCATCGTGTGCGAGAACCAATGAGTTGACATGGCCAACGTTTCTTGAAGATTTTAAACGTGCCATTTGCTCGCTGGAGGTTGATGCCGGTATCGGCATCCCTTACATAGCCTACGGTCTCCCCACCCATCGTGGTTGGGTTGAGAATCCGTCTCTGCTACCAGTTCTAGCTCAACTCACATTTGCTCGTCTACAGAAGTTAGCGAATGTGAGTTTTGAAGCTTTGTCTGCAGAGGAGCTTGTAAGGTGTGGTCTTTGCGACCCTATCCGTCTTTTTATTAAGATGGAGCCTCATAAGCAGAGCAAACTCGATGAAGGTCGCTACCGCCTCATCATGAGTGTCTCATTGGTTGATCAATTGGTAGCCCGGGTTTTATTTCAAAATCAGAATCAGCGGGAGATCGCCTTGTGGAGGGCGATTCCAAGCAAACCCGGATTTGGACTATCCACTGACGAGCAAGTCGCGGACTTTCTTGCTTGTCTTGCTAAGGTAGTGGGAACTACACCTACTGAAGCTTGCAGCTGTTGGCGCGAGCTTATGATTCCCACAGACTGCTCCGGTTTCGACTGGAGCGTCGCCGCTTGGATGCTCGAGGATGAAATGGAGGTGAGAAATCGCCTGACCATCAATAATACCGAGCTGTGTCGCCGCCTTCGGGCTGGCTGGACGCATTGCATCGCTAATTCTGTGCTTTGCTTGTCCGACGGCACCTTACTCGCCCAGACGGTGCCTGGCGTGCAGAAGTCCGGGAGTTACAATACTTCTTCATCTAACTCCCGAATTCGCGTCATGGCAGCCTTTCACGCAGGGGCTTCCTGGGCGATGGCGATGGGCGATGATGCCCTCGAGTCCACAGACAGCCAAATCCCTGTCTATAAAAATCTTGGATTTAAAGTCGAGGTCGCGGAACAGTTGGAATTTTGTTCACATGTTTTTGTGCGTCCGGACCTCGCCATTCCGACCGGAACCAATAAGATGCTGTATAAGCTTCTATTTGGTTACGATCCGGAATGTGGGAACCTAGAGGTACTCTCCAATTACATTGCTGCATGTTGGAGTATCCTCAACGAGCTTCGATCTGATCCCGAACTCGTTTCCCGCCTCTACTCGTGGCTGATTCCAACGCAGTCACAAAAGAATACCGCGGAGTAGAGCAAATAGGAAAACACATAGCCTCACATACACAGTTGCAAGTGGAGGAGTGTTTAGTCTGTTACCACGCGACGTTAAACAATTGATTTTAAGTTTATTGCCGGTTTTGGGCTTGGCTTTTTAGCCGCCATACCGATTTCAGTCGTCGCGATTTATTTTATCTACCTAAAGATCTCAGCCAACGTACGCTCAATTGTTAATGAATACGGGCGGGGCTAGATCTAGAAATGGACGTCGTAGAGTTCGACTACCACGCCGCCTCCAGCGGCGTAGGCCAGCTCAGCCAATCGTTGTGGTCTCGGGACCCAACCAGATTCGACGCCGTCGACGACGAAGAGGAAATTCTTCACGTCGAGGAGCAAATGCAGCTCCAGGAAGAGGAGGCTCTCGTGAAACATTTGTTTTCACGAAGGATGACCTCGCGGGCAACACCTCTGGAAGTCTCACCTTCGGGCCGAGTCTTTCAGAGTGTCCAGCATTCGCGACAGGAATTCTCAAAGCCTACCATGAGTATCGTATCTCACAGTGTACTTTGGAGTTCATCTCCGAAGCCCCTTCCACGGCGTCCGGTTCAATCGCTTATGAGCTGGATGCACACTGCAAAATCTCCTCCCTCTCCTCAAAAATCAACAAGTTTGGAATCACTAAAGGAGGGAAGAGGAGCTTTGCGGCGTCCAAAATCAACGGGATAGAGTGGCATGATTCTTCTGAGGATCAGTTTCGCCTTTTATACAAAGGCAATGGCGCTTCCAATATCGCTGGCTCTTTTCGAGTCACCATTGTCGTCGCCACTCAAAACCCGAAATAGGTAGACGGCGAACCAGGGCCCAGTCCAGGGCCAGACCCCCCCCCTCCAGTACCAGCTCCCCAACCACACAAGCACGAACGTTTCATTGCTTATGTTGGTATCCCTATGCTTACAATTCAAGCGAAGGAGAATGACGACGGGATCATTCTACGATCACTCGGACCACAACGCATGAAGTATATTGAGGATGAAAATCAAAATTATACAAATATTGATTCTCAATATTACTCTCAAACCAGTGTCTCCGCTGTACCCATGTACTACTTCAATGTCCCTTCGGGCAAGTGGAGTGTTGATATCAGTTGTGAAGGTTATCAGTGCACTTCAAGCACAACTGATCCCAACCGTGGACGAAGCGATGGTTTGATAGCGTATTCAAATAGTTCCGATGATTATTGGAATGTGGGAGAAGCTGATGGTGTTACAATCACCAAGTTGACCAATGACAACACGTGCAGGATGGGTCACCCCGATCTCGAGGTGAATTCCTGTCATTTTCGTGACGGTCAACTGTTGGAGCGTGACGCCACTATTTCCTTTCACGTTGAGGCTTCCAGTGATGGAAGATTCTTCTTGATCGGTCCCGCCATTCAGAAAATGGAAAAGTATAATTATACGATTTCGTATGGTGAGTGGACCGATCGTGATATGGAATTGGGTCTAATCACAGTTATTCTTGATGAACATCTTGAGGATTCTGGTTCGCGATTTGGTAGGATCCAACGGAGGAAGACACGAGATGGACACGTGCATCTTACCTCCATAGCGGAAAGACAACCGCTTACACAAACTCCTCGTTCTTTGCGAGTTCTCGATACTGGTTCGAGACATAAGTCCCTCTCACCTGAGAGGGCACGATCCCCCTCCCCCCTTAGGGAACCTTCACCACCTCCTCCGGAAAATCCGGAGATATGGCAAGTGGATTGGCATACTCCGAAATCTGGAGAACCTTTTGCTCCCTTGCCCCAGGATTTCAAACGTCCTGAGAGTGCGTTGAAGGGTGTCTTGCGTGAAGCTCGCGGTAACACCGGAAATTATGAGCGAGATTACGCGGATAAGGAAAATTGGGATCCTTCAACTCTGGAGGACCCCACTCGCTTTGGCACTAACTGGACTTCTTCCCAGATTGCTGAAGCGCAAGAGCAGACGGATGCTAAACGCAAAACTCGTTTCTCTCTTGTTCCTCGTTCTTTTAAGGGAGGTTCCTCCTTAGGTGGAGGGAGCCTTACGGGAGGTTCTCTGCGTAATACCCTTCGGGGTCGCTTAGAGAAGTTGACTGCTGAGCAGCTGATCCAATATCATAGGATCAAGCAGCAGCAGGGATCTTCCGTCGCCCAGATGTTCCTTACGGATTGCCTGGGTGACGGTTAACTTGCTCGTTTATTAATGAGCCCACACTGCAAGTAGCTTAGGCGAGTGCGTGTGGTTCCAAGTTTGGTCTACAATAAAGACTGGTCTTTTGGAGACGACCATAAATACATCTCCCCCAGTAAGTGTTCTGGGTTATCAAAAACCCGGGCAGTTTGTGGTCTGCCTATCAAAACCAACAGAAGGGTAGCAACCTTCCCGCCCCACTCGTTGTCTGCAGCGAGTGAGGGTTCCATTAGCTGGAGAGCAAACTTAGAGGGTAATTCCTCTTTGTTTGTTT

#### >AspPolV-1-NL1-14

TTCAGACGGTGTGCTCTTCCGATCCGAGAGAGCAGCTTTGAGCGCCTCCAACCCCCTTCCAAACCTTCAACTTAACCGTTTAAAAGTTGTATGTTTGTGATAACCTCTGAAGGATTCTTCTTGATAGACTCTTCTATCACTCCTTCTTTTGAGTTGTTGATTCTTCTTCTCCGCTCACTTGTTGCGTTCTGCATTTCTGCCCTACGCACCTCCTCGGTACCTTACGATGTCTTCTTCTTCAACTTTCATCGTTCTTTTCTTTTTGTTCTCCCTCTCCTCTGTGGTGTCCCCGTCCGCAACCTCTGCGACGGGCGATTTCGCGCCGCTCTCGACCTTCTCCCCTCTTTTCTGGAATGGGGCATTGTTACCCAATGCCACCCTCCTTTCCAGACTACCCGACGCCATGTCATTTTCGACATGCGCTTGTCCGGATCCGCCTCTCAATACATCACTCACTTACAACGAGTTGATGCGGATCTTCTGGGAAAAGGTCTCCTCCGACACCCAGAGCTTCTCCTCTCGCGCAATGGCAACCCTCAGCGAATACTCTGTTGCCTTGCTCGCCTCCGCTCGCAAGCACGCCTCCAATGCGCTCGAGTCCTTACTGTGGATTCCCGTCGCACTTTTTTGGACCAGCTCGTACTACGCGGGGGTAGTGGTGTGGGTTTGCTTGACCCAATATACTGTGATCGCGTTGGCGTATTTGTCGGTCGGTTGCTTCATAGCACTGGCTTGGAACGCCGCGATCTGGGTTTGTTCACGCTTTCCTATATTCCTTCTCTGTACACCCTTTTACGTCCTGAGGAGTGTTTGGAAGATTCTGTCATCCAAAAGCTTCTCCGTGAAGGTTGTCAATGAGAAAGCCGTGGACGGGTATGTGGATTATTCCATACCCCAGGACCCACCGAGAGGATCGACTGTACGCCTTCTTTATCCGAATGGCAACCCTTTGGGGTATGCTACTTGCGTACGGTTGTTCAATGGGGAGAACGCCCTCATAACCAGCTATCACTGCGTCGTTGAGGAAGGTGCTCTAGTTCACTCCACTCGCACCGGCAACAAATTGCCCCTTTCCCTGTTCAAACCCCTTTATGAGGATCAACGTGGTGATCTTTCGATCATGTGTGGTCCCCCTAATTGGGAAGGTTTGATGGGATGCAAGGGAGTTCATGCCGTGACTTGCGATCGCTTGGGACGCGGACCAGCCACTTTCTTCGTTTACGATAATAGCTGGAAAGCGAGGTCAGCCCAGATCACTGGTCGCTATGAGGATTTCGCTCAGGTTCTTTCTAATACGGAACCTGGAGTGAGTGGGGCAGGGTATTTTAATGGCAAAACCCTGCTCGGTGTGCATAAGGGACACACCGACGTGGCTGAACACAACTTCAATTTAATGTCGCCACTCCCATGTCTCCCTGGTCTGACAGCCCCCATATACGTTTATGAGACCACAAACGTGCAGGGGAGAATCTTCGACGAGGACTTCGTTATACAATTTAAGAAGGGAAGCTGGTTCGATCAGATGATCGAACACGCTGAGAACCACGACGTGCTACCACCTTTGGTACGGCGTCCCGATGGTTCTCTCCGCTTTGCGGATGAGAAATCCCTCCCAATCGAGTTCTCGGGAAACGGAGCAGGCAGCGCCGTCTGCCCACCAAACGAGCCTGTCAAGACCCCGTCAGCCCCCCCCCCGATGCCCTCCCCCTCCCTTCTAGGAAGCTTTGTCGCTCCGTCCTCCGACACCCCTGTGACCGTCGGCGTTTCTTCGAACGCCTTGCAGGAAGGAGTCTTAAAGGCGTTGGTAGCGAAGTTCGACTTCCAGAAGGTAGAGCAGAATGTCGCAGAATTAATCGCTGCGAAAATGCTCTCCCGTCCTCAGCGAACGCGTGGTACCCGAGGGAGGAAGCGAAGCAATTCCAAAACTACTTCGCCTCCCTCTACAAGTGGTCAGAGTCAGCCCCCTCCGAAGCCCCAGGCTTCAAAACCGTCGGCAGGCTCCCCTCGTACTACCACCCCTCCCCAAAACGGGAAAGCAAATGGGGGAAAAAGCTCACCCGGCAATATCCCGAGCTGGGTGAGAAAACCACAGGCTTCGGCTGGCCGGGGGCTGGCGCAGAAGCGGAGCTGAAATCCCTTCAGCTCCAAGCGGCCAGGTGGCTCCAACGCGCGGAGACCGCCATCATTCCAAATCAGGCCGATCGGGAGCGCGTAATCGCTCGCACCGTTGACGCGTACTGTGCCGTTCATACGGAAGTACCATCGTGTGCGAGAACCAATGAGTTGACATGGCCAACGTTTCTTGAAGATTTTAAACGTGCCATTTGCTCGCTGGAGGTTGATGCCGGTATCGGCATCCCTTACATAGCCTACGGTCTCCCCACCCATCGTGGTTGGGTTGAGAATCCGTCTCTGCTACCAGTTC TAGCTCAACTCACATTTGCTCGTCTACAGAAGTTAGCGAATGTGAGTTTTGAAGCTTTGTCTGCAGAGGAGCTTGTAAGGTGTGGTCTTTGCGACCCTATCCGTCTTTTTATTAAGATGGAGCCTCATAAGCAGAGCAAACTCGATGAAGGTCGCTACCGCCTCATCATGAGTGTCTCATTGGTTGATCAATTGGTAGCCCGGGTTTTATTTCAAAATCAGAATCAGCGGGAGATCGCCTTGTGGAGGGTGATTCCAAGCAAACCCGGATTTGGACTATCCACTGACGAGCAAGTCGCGGACTTTCTTGCTTGTCTTGCTAAGGTAGTGGGAACTACACCTACTGAAGCTTGCAGCTGTTGGCGCGAGCTTATGATTCCCACAGACTGCTCCGGTTTCGACTGGAGCGTCGCCGCTTGGATGCTCGAGGATGAAATGGAGGTGAGAAATCGCCTGACCATCAATAATACCGAGCTGTGTCGCCGCCTTCGGGCTGGCTGGACGCATTGCATCGCTAATTCTGTGCTTTGCTTGTCCGACGGCACCTTACTCGCCCAGACGGTGCCTGGCGTGCAGAAGTCCGGGAGTTACAATACTTCTTCATCTAACTCCCGAATTCGCGTCATGGCAGCCTTTCACGCAGGGGCTTCCTGGGCGATGGCGATGGGCGATGATGCCCTCGAGTCCACAGACAGCCAAATCCCTGTCTATAAAAATCTTGGATTTAAAGTCGAGGTCGCGGAACAGTTGGAATTTTGTTCACATGTTTTTGTGCGTCCGGACCTCGCCATTCCGACCGGAACCAATAAGATGCTGTATAAGCTTCTATTTGGTTACGATCCGGAATGTGGGAACCTAGAGGTACTCTCCAATTACATTGCTGCATGTTGGAGTATCCTCAACGAGCTTCGATCTGATCCCGAACTCGTTTCCCGTCTCTACTCGTGGCTGATTCCAACGCAGTCACAAAAGAATACCGCGGAGTAGAGCAAATAGGAAAACACATAGCCTCACATACACAGTTGCAAGTGGAGGAGTGTTTAGTCTGTTACCACGCGACGTTAAACAATTGATTTTAAGTTTATTGCCGGTTTTGGGCTTGGCTTTTTAGCTGCCATACCGATTTCAGTCGTCGCGATTTATTTTATCTACCTAAAGATCTCAGCCAACGTACGCTCAATTGTTAATGAATACGGGCGGGGCTAGATCTAGAAATGGACGTCGTAGAGTTCGACTACCACGCCGCCTCCAGCGGCGTAGGCCAGCTCAGCCAATCGTTGTGGTCTCGGGACCCAACCAGATTCGACGCCGTCGACGACGAAGAGGAAATTCTTCACGTCGAGGAGCAAATGCAGCTCCAGGAAGAGGAGGCTCTCGTGAAACATTTGTTTTCACGAAGGATGACCTCGCGGGCAACACCTCTGGAAGTCTCACCTTCGGGCCGAGTCTTTCAGAGTGTCCAGCATTCGCGACAGGAATTCTCAAAGCCTACCATGAGTATCGTATCTCACAGTGTACTTTGGAGTTCATCTCCGAAGCCCCTTCCACGGCGTCCGGTTCAATCGCTTATGAGCTGGATGCACACTGCAAAATCTCCTCTCTCTCCTCAAAAATCAACAAGTTTGGAATCACTAAAGGAGGGAAGAGGAGCTTTGCGGCGTCCAAAATCAACGGGATAGAGTGGCATGATTCTTCTGAGGATCAGTTTCGCCTTTTATACAAAGGCAATGGCGCTTCCAATATCGCTGGCTCTTTTCGAGTCACCATTGTCGTCGCCACTCAAAACCCGAAATAGGTAGACGGCGAACCAGGGCCCAGTCCAGGGCCAGACCCCCCCCCTCCAGCACCAGCTCCCCAACCACACAAGCACGAACGTTTCATTGCTTATGTTGGTATCCCTATGCTTACAATTCAAGCGAAGGAGAATGACGACGGGATCATTCTACGATCACTCGGACCACAACGCATGAAGTATATTGAGGATGAAAATCAAAATTATACAAATATTGATTCTCAATATTACTCTCAAACCAGTGTCTCCGCTGTACCCATGTACTACTTCAATGTCCCTTCGGGCAAGTGGAGTGTTGATATCAGTTGTGAAGGTTATCAGTGCACTTCAAGCACAACTGATCCCAACCGTGGACGAAGCGATGGTTTGATAGCGTATTCAAATAGTTCCGATGATTATTGGAATGTGGGAGAAGCTGATGGTGTTATAATCACCAAGTTGACCAATGACAACACGTACAGGATGGGTCACCCCGATCTCGAGGTGAATTCCTGTCATTTTCGTGACGGTCAACTGTTGGAGCGTGACGCCACTATTTCCTTTCACGTTGAGGCTTCCAGTGATGGAAGATTCTTCTTGATCGGTCCCGCCATTCAGAAAATGGAAAAGTATAATTATACGATTTCGTATGGTGAGTGGACCGATCGTGATATGGAATTGGGTCTAATCACAGTTATTCTTGATGAACATCTTGAGGATTCTGGTTCGCGATTTGGTAGGATCCAACGGAGGAAGACACGAGATGGACACGTGCATCTTACCTCCATAGCGGAAAGACAACCGCTTACACAAACTCCTCGTTCTTTGCGAGTTCTCGATACTGGTTCGAGACATAAGTCCCTCTCACCTGAGAGGGCACGATCCCCCTCCCCCCTTAGGGAACCTTCACCACCTCCTCCGGAAAATCCGGAGATGTGGCAAGTGGATTGGCATGCTCCGAAATCTGGAGAACCTTTTGCTCCCTTGCCCCAGGATTTCAAACGTCCTGAGAGTGCGTTGAAGGGTATCTTGCGTGAAGCTCGCGGTAACACCGGAAATTATGAGCGAGATTACGCGGATAAGGAAAATTGGGATCCTTCAACTCTGGAGGACCCCACTCGCTTTGGCACTAACTGGACTTCTTCCCAGATTGCTGAAGCGCAAGAGCAGACGGATGCTAAACGCAAAACTCGTTTCTCTCTTGTTCCTCGTTCTTTTAAGGGAGGTTCCTCCTTAGGTGGAGGGAGCCTTACGGGAGGTTCTCTGCGTAATACCCTTCGGGGTCGCTTAGAGAAGTTGACTGCTGAGCAGCTGATCCAATATCATAGGATCAAGCAGCAACAGGGATCTACCGTCGCCCAGATGTTCCTTACGGATTGCCTGGGTGACGGTTAACTTGCTCGTTTATTAATGAGCCCACACTGCAAGTAGCTTAGGCGAGTGCGTGTGGTTCCAAGTTTGGTCTACAATAAAGACTGGTCTTTCGGAGACGACCATAAATACATCTCCCCCAGTAAGTGTTCTGGGTTATCAAAAACCCGGGCAGTTTGTGGTCTGCCTATCAAAACCAACAGAAGGGTAGCAACCTTCCCGCCCCACTCGTTGTCTGCAGCGAGTGAGGGTTCCATTAGCTGGAGAGCAAACTTAGAGGGTAATTCCTCTTTGTTTGTTT

#### >AspPolV-2-NL1-24a

TTCAGACGGTGTGCTCTTCCGAACCGAGAGAGCAGCTTTGAGCGCCTCCAACTCCTTTCCAAACCTTCAACTTAACCGTCTAAAAGTTGTATGTTTGTGATTACCCCTGAAGGAATTTTCCTGATAGATTCTTCTATCTCTCCTTCCGTTGAGTTGTTAACCCTCCTCCTTCGCTCTTTGATCGCGTTCTGCTTTTCTGCTCAACGCACCTCCTCGGTTCCCGACGATGTCTTCTTCTACAATTTCCGTCGTTCTTTTCTTTTTGTTCTCCCTCTTCTTTGTGGCGTCCCCGTCCGCTACTTCAGCGACGGGCGCTTCCGCGCCACTCTCAACCTTCTCCCCGCTTTTCTGGAATGGGGCATTGTTACCCAATGCCACCCTCCTTTCCAGATTACCAAACGACATGTCATCTTCGACATGCGTTTGTTCGGATCCGCCTCTCGTTATATCACTCACTTACAACGAGTTGATGCGGATCTTCTCGGAGAAGGTCTGCTCCGACACTCGGAGCTTCTCCTCTCGCGCAATGGCAACCCTCAGCGAACACTCTGTTGCCTTGCTTACCTCCGCGCACAAGCACGCTTCCAGAGCGCTCGAGTCCTTTCTGTGGATTCCTGTCGCGCTTTTCTGGACCAGCTTGTATTACGTGGGGGTAGTAGTGTGGGTCTGCTTGACCCAGTACACTGCAATCGCATTGGCGTACTTATTGGTCGGTTGCTTCATAGCACTGGTTTGGAACGCCGCGATTTGGGTTTGTTCACGCTTTCCTTTATTCCTTCTTTGTACACCCTTCTACGTCCTGAAGAGTGTTTGGAAGACTCTGTCATTCAAACGCTCTTCCGTGAAGGTTGTGAATGAGAAAGCAGTGGACGGGTATGTGGATTATTCCATACCCCAAGACCCACCGAGAGGATCAACCGTGCGCCTTTTATACCCGAATGGAAATCCACTGGGGTATGCTACTTGCGTTCGGTTGTTTAATGGGGAGAACGCCCTCATAACCAGCTATCACTGCGTTATTGAGGAAGGTGTTCTGGTTCACTCCACTCGCACCGGCAACAAATTACCCCTTTCCCTTTTCAAGCCCCTTTATGAGGACCAACGTGGTGATCTTTCGATTATGTGCGGTCCTCCTAACTGGGAAGGTTTGATGGGATGCAAGGGAGTGCATGCCGTGACTTGCGATCGTTTGGGTCGTGGACCAGCTACCTTCTTTGTTTATGACAATAGCTGGAAGGCGAGGTCAGCCCAGATCGCTGGTCGCTATGAGGATTTCGCTCAGGTTCTTTCTAACACCGAACCTGGAGTGAGTGGGGCAGGGTACTTTAATGGTAAAACCCTGCTCGGTGTGCATAAGGGACACACCGACGTGGCTGAACACAATTTCAACTTGATGTCGCCACTCCCATGTCTCCCTGGTCTGACAGCCCCCATATACGTTTATGAAACCACGAACGTACAGGGGAGAATCTTCGACGAGGACTTTGTGGTCCAATTCAAGAAGGGAAGCTGGTTCGATCAGATGATCGAACACGCTGAGAACCACGATGTGCTACCTCCTTTGGTGCGCCGTCCTGATGGCTCTCTCCGCTTTGCGGATGAAAAATCCCTTCCCGCCAAATTCTCGGGAAACGAAGCTGGCAGCGCCGTCTGCCCACCAAACGAGCCTGTCAAGACCCCGTCAGCCCCCCCCCCGATGCCCTCCCCCTCCCTTCTAGGAAGCTTTGTCGCTCCACCCTCCGACACCCCTGCGACCGTCGGCGTTTCTTCAAACGCCTTGCAGGAAGGTGTCCTCAAGGCGTTGGTAGCGAAGTTCGACTTCCAGAAGGTCGAGCAGAATGTCGCAGAATTGATTGCTGCGAAAATGCTCTCCCGTCCTCAGCGAACGCGTGGTACCCGAGGGAGGAGACGAAGCAATTCCAAAACTACTTCGCCTCCCTCTACAAGTGGTCAGAGCCAGCCCCCTCTGAAGCCCCAGGCTTCAAAACCGTTGGCAGGCTCCCCTCGTACTACCACCCCTCCCCAAAACGAGAGAGCAAGTGGGGGAAAAAGCTCACCCGGCAACATCCCGAGCTGGGTGAGAAAACCCAAGGCTTCGGCTGGCCAGGGGCCGGCGCAGAAGCGGAGTTGAAATCTCTTCAGCTCCAAGCGGCCAGGTGGCTCCAACGCGCGGAGACCGCCACCATTCCAAATCAGGCCGATCGGGAGCGCGTAATCGCTCGCACCGTTGACGCGTACTGTGCCGTTCATACGGAAGTACCATCGTGTGCGAGAACCAATGAGTTGACATGGCCAACGTTTCTTGAAGATTTTAAACGTGCCATTTGCTCGCTGGAGGTTGATGCCGGTATCGGCATCCCTTACATAGCCTACGGTCTCCCCACCCATCGTGGTTGGGTTGAGAATCCGTCTCTGCTACCAGTTCTAGCTCAACTCACATTTGCTCGTCTACAGAAGTTAGCGAATGTGAGTTTTGAAGCTTTGTCTGCAGAGGAGCTTGTAAGGTGTGGTCTTTGCGACCCTATCCGTCTTTTTATTAAGATGGAGCCTCATAAGCAGAGCAAACTCGATGAAGGTCGCTACCGCCTCATCATGAGTGTCTCATTGGTTGATCAATTGGTAGCCCGGGTTTTATTTCAAAATCAGAATCAGCGGGAGATCGCCTTGTGGAGGGCGATTCCAAGCAAACCCGGATTTGGACTATCCACTGACGAGCAAGTCGCGGACTTTCTTGCTTGTCTTGCTAAGGTAGTGGGAACTACACCTACTGAAGCTTGCAGCCGTTGGCGCGAGCTTATGATTCCCACAGACTGCTCCGGTTTCGACTGGAGCGTCGCCGCTTGGATGCTCGAGGATGAAATGGAGGTGAGAAATCGCCTGACCATCAATAATACCGAGCTGTGTCGCCGCCTTCGGGCTGGCTGGACGCATTGCATCGCTAATTCTGTGCTTTGCTTGTCCGACGGCACCTTACTCGCCCAGACGGTGCCTGGCGTGCAGAAGTCCGGGAGTTACAATACTTCTTCATCTAACTCCCGAATTCGCGTCATGGCAGCCTTTCACGCAGGGGCTTCCTGGGCGATGGCGATGGGCGATGATGCCCTCGAGTCCACAGACAGCCAAATCCCTGTCTATAAAAATCTTGGATTTAAAGTCGAGGTCGCGGAACAGTTGGAATTCTGTTCACACATTTTTGTGCGTCCGGACCTCGCCATTCCGACCGGGACCAACAAAATGCTGTACAAGCTTTTGTTTGGTTACGATCCGGAATGTGGGAACCTAGAGGTACTCTCCAACTACATTGCTGCATGTTGGAGCATCCTCAACGAGCTTCGATCTGATCCCGAACTCGTTTCCCGCCTCTACTCGTGGCTGATTCCAACGCAGTCACAAAAGAATACCGCGGAGTAGAGCAAATAGGAAAACACATAGCCTCACATACACAGTTGCAAGTGGAGGAGTGTTTTAGACTGTTACCACGCGACGTTAAACAATTGATTTTAAGTTTATTGCCGGTTTTGGGCTTGGCTTTTTAGCCGCCATACCGATTTCAGTCGTCGCGATTTATTTTATCTACCTAAAGATCTCAGCCAACGTACGCTCAATTGTTAATGAATACGGGCGGGGCTAGATCTAGAAATGGACGTCGTAGAGTTCGATTACCACGCCGCCTCCAGCGGCGTAGG CCAGCTCAGCCAATCGTTGTGGTCTCAGGACCCAACCAGATTCGACGCCGTCGACGACGAAGAGGAAATTCTTCACGTCGAGGAGCAAATGCAGCTTCAGGAAGAGGAGGCTCTCGTGAAACATTCGTTTTCACGAAGGACGACCTCGCGGGCAACACCTCTGGAAGTCTCACCTTCGGGCCGAGTCTTTCAGAGTGTCCAGCATTCGCGACAGGAATTCTCAAAGCCTACCATGAGTATCGTATCTCACAGTGTACTTTGGAGTTCATCTCCGAAGCCCCTTCCACGGCGTCCGGTTCAATCGCTTATGAGCTGGATGCACATTGCAAAATCTCCTCTCTCTCCTCGAAAATCAACAAGTTTGGAATCACCAAAGGAGGGAAGAGGAGCTTTGCGGCGTCCAAAATCAACGGGATAGAGTGGCATGATTCTTCCGAGGATCAATTTCGCCTTTTGTATAAAGGCAATGGCGCTTCTAATATTGCTGGCTCTTTTCGAGTCACTATTGTCGTCGCCACTCAAAACCCGAAATAGGTAGACGGCGAACCAGGGCCCAGCCCAGGGCCAGACCCCCCCCCTCCAGCACCAGCTCCCCAGCCACGTAAACATGAACGTTTCATTGCTTACGTTGGCATCCCTATGCTTACAATTCAAGCAAAGGAGAATGACGATGGGATCATTCTAAGGTCACTCGGACCCCAACGCATGAAGTATATTGAGGATGAAAATCAAAATTACACTAATATTGATTCCCAATACTACTCTCAAACCAATGTGTCCGCTGTCCCCATGTACTACTTCAATGTCCCAGCGGGCAAGTGGAGTGTTGACGTCAGTTGTGAAGGTTATCAGTGCACTTCAAGCACGACTGACCCCAACCGTGGACGAAGCGATGGTTTGATTGCGTATTCAGACAACTCCGATGACTATTGGAATGTGGGTGAAGCTGATGGTGTGATAATCACCAAGTTGACCAATGACAACACGTACAGGATGGGTCACCCCGATCTCGAGGTGAATTCCTGTCATTTTCGTGATGGTCAGTTGCTGGAGCGTGATGCCACGATTTCTTTTCACGTTGAGGCTTCCAGTGATGGAAGATTCTTCTTGATTGGTCCCGCCATTCAGAAAATGGAAAAGTATAATTATACGATTTCGTATGGTGAGTGGACCGACCGTGATATGGAATTGGGTCTAATCACAGTTATTCTTGACGAACATCTTGAAGATTCTGGTTCGCGATTTGGTAAGATCCAACGGAGGAAGACACGAGATGGGCACGTGCGCCTTACCTCCGTAGCGGAAAGACAACCGCTTGTACAAACTCCTCGCCCCTTGCGAGTTCTCGATACTGGTTCGAGACGTAAGTCCCTCTCCCCGGAGAGGGCAAACTCCCCCCCCCCCCTTTGAGAACCGTCACCACCTCCTCCGGAAGATCCAGAGTTGGGGCAGGTCGGTTGGTACACCCCAAAATCTGGAGAACCACTGGCCCCTTTGCCCCAGGACTTTAAACGTCCTGAGAGTGTGTTGAAGGGTATCCAGCGTGAGTCCCGTGGTAACACCGGTAACTATGAAAAGGACTACGCGGATAAGGAAAATAGGGATCCTTCAACTTTGGAAGACCCAACTCGCTTCGGAACGAACTGGACAAGCTCCCAGATCGCCGAAGCACAAGAGCAGACGGATGCTAAACGCAAAACTCGTTTCTCTCTTGTTCCTCGTTCCTTTAAGGGAGGTTCCTCCATTAGTGGGGGAAGCTTATCGGGGGGTTCTCTTCGAAGCACTCTACGGAGTCGCTTGGAGAAGCTGACTGCTGAGCAGCAAATCCAATATCATCGGATTAAGCAGCAGCAGGGAGCTACCACCGCTCAGATGTTCCTTTCGGATTGCCTGGGTGATGGTTAACTTGCTCGTTTAATAACGAGCCCACACTGCTAGTAGCCTAGGCGAGCGCGTGTGGTTCCAAGTTTGGTCAACAATAAAGACTGGTCCTTCGGAGACGACCTTAAATACATCTCCCCCAGTAAGTGTTCTGGGTTATCAAAACCGGGGCAGTTCGTGGTCTGCCGATCAAAACCAACAGAAGGGTAGCAACCTTCCCACCCCCTGACACGTATTGTGTTAGTTGAGGGTTCCGTTAGCTTAGGAGAGCAAACCGCGAGGGTCATTCCTCGAGGTTTGTTCG

#### >AspPolV-2-NL1-24b

TTCAGACGGTGTGCTCTTCCGAACCGAGAGAGCAGCTTTGAGCGCCTCCAACTCCTTTCCAAACCTTCAACTTAACCGTCTAAAAGTTGTATGTTTGTGATTACCCCTGAAGGAATTTTCCTGATAGATTCTTCTATCTCTCCTTCCGTTGAGTTGTTAACCCTCCTCCTTCGCTCTTTGATCGCGTTCTGCTTTTCTGCTCAACGCACCTCCTCGGTTCCCGACGATGTCTTCTTCTACAATTTCCGTCGTTCTTTTCTTTTTGTTCTCCCTCTTCTTTGTGGCGTCCCCGTCCGCTACTTCAGCGACGGGCGCTTCCGCGCCACTCTCAACCTTCTCCCCGCTTTTCTGGAATGGGGCATTGTTACCCAATGCCACCCTCCTTTCCAGATTACCAAACGACATGTCATCTTCGACATGCGTTTGTTCGGATCCGCCTCTCGTTATATCACTCACTTACAACGAGTTGATGCGGATCTTCTCGGAGAAGGTCTGCTCCGACACTCGGAGCTTCTCCTCTCGCGCAATGGCAACCCTCAGCGAACACTCTGTTGCCTTGCTTACCTCCGCGCACAAGCACGCTTCCAGAGCGCTCGAGTCCTTTCTGTGGATTCCTGTCGCGCTTTTCTGGACCAGCTTGTATTACGTGGGGGTAGTAGTGTGGGTCTGCTTGACCCAGTACACTGCAATCGCATTGGCGTACTTATTGGTCGGTTGCTTCATAGCACTGGTTTGGAACGCCGCGATTTGGGTTTGTTCACGCTTTCCTTTATTCCTTCTTTGTACACCCTTCTACGTCCTGAAGAGTGTTTGGAAGACTCTGTCATTCAAACGCTCTTCCGTGAAGGTTGTGAATGAGAAAGCAGTGGACGGGTATGTGGATTATTCCATACCCCAAGACCCACCGAGAGGATCAACCGTGCGCCTTTTATACCCGAATGGAAATCCACTGGGGTATGCTACTTGCGTTCGGTTGTTTAATGGGGAGAACGCCCTCATAACCAGCTATCACTGCGTTATTGAGGAAGGTGTTCTGGTTCACTCCACTCGCACCGGCAACAAATTACCCCTTTCCCTTTTCAAGCCCCTTTATGAGGACCAACGTGGTGATCTTTCGATTATGTGCGGTCCTCCTAACTGGGAAGGTTTGATGGGATGCAAGGGAGTGCATGCCGTGACTTGCGATCGTTTGGGTCGTGGACCAGCTACCTTCTTTGTTTATGACAATAGCTGGAAGGCGAGGTCAGCCCAGATCGCTGGTCGCTATGAGGATTTCGCTCAGGTTCTTTCTAACACCGAACCTGGAGTGAGTGGGGCAGGGTACTTTAATGGTAAAACCCTGCTCGGTGTGCATAAGGGACACACCGACGTGGCTGAACACAATTTCAACTTGATGTCGCCACTCCCATGTCTCCCTGGTCTGACAGCCCCCATATACGTTTATGAAACCACGAACGTACAGGGGAGAATCTTCGACGAGGACTTCGTTACACAATTTAAGAAGGGAAGCTGGTTCGATCAGATGATCGAACACGCTGAGAACCACGACGTGCTACCACCTTTGGTACGGCGTCCCGATGGTTCTCTCCGCTTTGCGGATGAGAAATCCCTCCCAATCGAGTTCTCGGGAAACGGAGCAGGCAGCGCCGTCTGCCCACCAAACGAGCCTGCCGAGACCCCGTCAGCCCCCCCCCCGATGCCCTCCCCCTCCCTTCTAGGAAGCTTTGTCGCTCCGTCCTCCGACACCCCTGTGACCGTCGGCGTTTCTTCGAACGCCTTGCAGGAAGGAGTCTTAAAGGCGTTGGTAGCGAAGTTCGACTTCCAGAAGGTAGAGCAGAATGTCGCAGAATTAATCGCTGCGAAAATGCTCTCCCGTCCTCAGCGAATGCGTGGTACCCGAGGGAGGAAGCGAAGCAATTCCAAAACTACTTCGCCTCCCTCTACAAGTGGTCAGAGTCAGCCCCCTCCGAAGCCCCAGGCTTCAAAACCGTCGGCAGGCTCCCCTCGTACTACCACCCCTCCCCAAAACGGGAAAGCAAATGGGGGAAAAAGCTCACCCGGCAATATCCCGAGCTGGGTGAGAAAACCACAGGCTTCGGCTGGCCGGGGGCTGGCGCAGAAGCGGAGCTGAAATCCCTTCAGCTCCAAGCGGCCAGGTGGCTCCAACGCGCGGAGACCGCCATCATTCCAAATCAGGCCGATCGGGAGCGCGTAATCGCTCGCACCGTTGACGCGTACTGTGCCGTTCATACGGAAGTACCATCGTGTGCGAGAACCAATGAGTTGACATGGCCAACGTTTCTTGAAGATTTTAAACGTGCCATTTGCTCGCTGGAGGTTGATGCCGGTATCGGCATCCCTTACATAGCCTACGGTCTCCCCACCCATCGTGGTTGGGTTGAGAATCCGTCTCTGCTACCAGTTCTAGCTCAACTCACATTTGCTCGTCTACAGAAGTTAGCGAATGTGAGTTTTGAAGCTTTGTCTGCAGAGGAGCTTGTAAGGTGTGGTCTTTGCGACCCTATCCGTCTTTTTATTAAGATGGAGCCTCATAAGCAGAGCAAACTCGATGAAGGTCGCTACCGCCTCATCATGAGTGTCTCATTGGTTGATCAATTGGTAGCCCGGGTTTTATTTCAAAATCAGAATCAGCGGGAGATCGCCTTGTGGAGGGCGATTCCAAGCAAACCCGGATTTGGACTATCCACTGACGAGCAAGTCGCGGACTTTCTTGCTTGTCTTGCTAAGGTAGTGGGAACTACACCTACTGAAGCTTGCAGCCGTTGGCGCGAGCTTATGATTCCCACAGACTGCTCCGGTTTCGACTGGAGCGTCGCCGCTTGGATGCTCGAGGATGAAATGGAGGTGAGAAATCGCCTGACCATCAATAATACCGAGCTGTGTCGCCGCCTTCGGGCTGGCTGGACGCATTGCATCGCTAATTCTGTGCTTTGCTTGTCCGACGGCACCTTACTCGCCCAGACGGTGCCTGGCGTGCAGAAGTCCGGGAGTTACAATACTTCTTCATCTAACTCCCGAATTCGCGTCATGGCAGCCTTTCACGCAGGGGCTTCCTGGGCGATGGCGATGGGCGATGATGCCCTCGAGTCCACAGACAGCCAAATCCCTGTCTATAAAAATCTTGGATTTAAAGTCGAGGTCGCGGAACAGTTGGAATTCTGTTCACACATTTTTGTGCGTCCGGACCTCGCCATTCCGACCGGGACCAACAAAATGCTGTACAAGCTTTTGTTTGGTTACGATCCGGAATGTGGGAACCTAGAGGTACTCTCCAACTACATTGCTGCATGTTGGAGCATCCTCAACGAGCTTCGATCTGATCCCGAACTCGTTTCCCGCCTCTACTCGTGGCTGATTCCAACGCAGTCACAAAAGAATACCGCGGAGTAGAGCAAATAGGAAAACACATAGCCTCACATACACAGTTGCAAGTGGAGGAGTGTTTTAGACTGTTACCACGCGACGTTAAACAATTGATTTTAAGTTTATTGCCGGTTTTGGGCTTGGCTTTTTAGCCGCCATACCGATTTCAGTCGTCGCGATTTATTTTATCTACCTAAAGATCTCAGCCAACGTACGCTCAATTGTTAATGAATACGGGCGGGGCTAGATCTAGAAATGGACGTCGTAGAGTTCGATTACCACGCCGCCTCCAGCGGCGTAGGCCAGCTCAGCCAATCGTTGTGGTCTCAGGACCCAACCAGATTCGACGCCGTCGACGACGAAGAGGAAATTCTTCACGTCGAGGAGCAAATGCAGCTTCAGGAAGAGGAGGCTCTCGTGAAACATTCGTTTTCACGAAGGACGACCTCGCGGGCAACACCTCTGGAAGTCTCACCTTCGGGCCGAGTCTTTCAGAGTGTCCAGCATTCGCGACAGGAATTCTCAAAGCCTACCATGAGTATCGTATCTCACAGTGTACTTTGGAGTTCATCTCCGAAGCCCCTTCCACGGCGTCCGGTTCAATCGCTTATGAGCTGGATGCACATTGCAAAATCTCCTCTCTCTCCTCGAAAATCAACAAGTTTGGAATCACCAAAGGAGGGAAGAGGAGCTTTGCGGCGTCCAAAATCAACGGGATAGAGTGGCATGATTCTTCCGAGGATCAATTTCGCCTTTTGTATAAAGGCAATGGCGCTTCTAATATTGCTGGCTCTTTTCGAGTCACTATTGTCGTCGCCACTCAAAACCCGAAATAGGTAGACGGCGAACCAGGGCCCAGCCCAGGGCCAGACCCCCCCCCTCCAGCACCAGCTCCCCAGCCACGTAAACATGAACGTTTCATTGCTTACGTTGGCATCCCTATGCTTACAATTCAAGCAAAGGAGAATGACGATGGGATCATTCTAAGGTCACTCGGACCCCAACGCATGAAGTATATTGAGGATGAAAATCAAAATTACACTAATATTGATTCCCAATACTACTCTCAAACCAATGTGTCCGCTGTCCCCATGTACTACTTCAATGTCCCAGCGGGCAAGTGGAGTGTTGACGTCAGTTGTGAAGGTTATCAGTGCACTTCAAGCACGACTGACCCCAACCGTGGACGAAGCGATGGTTTGATTGCGTATTCAGACAACTCCGATGACTATTGGAATGTGGGTGAAGCTGATGGTGTGATAATCACCAAGTTGACCAATGACAACACGTACAGGATGGGTCACCCCGATCTCGAGGTGAATTCCTGTCATTTTCGTGATGGTCAGTTGCTGGAGCGTGATGCCACGATTTCTTTTCACGTTGAGGCTTCCAGTGATGGAAGATTCTTCTTGATTGGTCCCGCCATTCAGAAAATGGAAAAGTATAATTATACGATTTCGTATGGTGAGTGGACCGACCGTGATATGGAATTGGGTCTAATCACAGTTATTCTTGACGAACATCTTGAAGATTCTGGTTCGCGATTTGGTAAGATCCAACGGA GGAAGACACGAGATGGGCACGTGCGCCTTACCTCCGTAGCGGAAAGACAACCGCTTGTACAAACTCCCCCCCCCTTGCGAGTTCTCGATACTGGTTCGAGACGTAAGTCCCTCTCCCCGGAGAGGGCAAACTCCCCCCCCCCCCGTAGAGAACCTTCACCACCTCCTCCGGAAGATCCAGAGTTGTGGCAGGGCGGTTGGTACACCCCAAAATCTGGAGAACCACTGGCCCCTTTGCCCCAGGACTTTAAACGTCCTGAGAGTGTGTTGAAGGGTATCCAGCGTGAGTCCCGTGGTAACACCGGTAACTATGAAAAGGACTACGCGGATAAGGAAAATAGGGATCCTTCAACTTTGGAAGACCCAACTCGCTTCGGAACGAACTGGACAAGCTCCCAGATCGCCGAAGCACAAGAGCAGACGGATGCTAAACGCAAAACTCGTTTCTCTCTTGTTCCTCGTTCCTTTAAGGGAGGTTCCTCCATTAGTGGGGGAAGCTTATCGGGGGGTTCTCTTCGAAGCACTCTACGGAGTCGCTTGGAGAAGCTGACTGCTGAGCAGCAAATCCAATATCATCGGATTAAGCAGCAGCAGGGAGCTACCACCGCTCAGATGTTCCTTTCGGATTGCCTGGGTGATGGTTAACTTGCTCGTTTAATAACGAGCCCACACTGCTAGTAGCCTAGGCGAGCGCGTGTGGTTCCAAGTTTGGTCAACAATAAAGACTGGTCCTTCGGAGACGACCTTAAATACATCTCCCCCAGTAAGTGTTCTGGGTTATCAAAACCGGGGCAGTTCGTGGTCTGCCGATCAAAACCAACAGAAGGGTAGCAACCTTCCCACCCCCTGACACGTATTGTGTTAGTTGAGGGTTCCGTTAGCTTAGGAGAGCAAACCGCGAGGGTCATTCCTCGAGGTTTGTTCG

### Supplemental File SF2. AspPolV virus protein sequences

#### >AspPolV-1-NL1-14_P0

MFVITSEGFFLIDSSITPSFELLILLLRSLVAFCISALRTSSVPYDVFFFNFHRSFLFVLPLLCGVPVRNLCDGRFRAALDLLPSFLEWGIVTQCHPPFQTTRRHVIFDMRLSGSASQYITHLQRVDADLLGKGLLRHPELLLSRNGNPQRILCCLARLRSQARLQCARVLTVDSRRTFLDQLVLRGGSGVGLLDPIYCDRVGVFVGRLLHSTGLERRDLGLFTLSYIPSLYTLLRPEECLEDSVIQKLLREGCQ

#### >AspPolV-1-NL1-2_P0

MFVITSEGFFLIDSSITPSFELLILLLRSLVAFCISALRTSSVPYDVFFFNFHRSFLFVLPLLCGVPVRNLCDGRFRAALDLLPSFLEWGTVTQCHPPFQTTRRHVIFDMRLSGSASQYITHLQRVDADLLGKGLLRHPELLLSRNGNPQRTLCCLARLRSQARLQCARVLTVDSRRTFLDQLVLRGGSGVGLLDPIYCDRVGVFVGRLLHSTGLERRDLGLFTLSYIPSLYTLLRPEECLEDSVIQKLLREGCQ

#### >AspPolV-1-NL1-3_P0

MFVITSEGFFLIDSSITPSFELLILLLRSLVAFCISALRTSSVPYDVFFFNFHRSFLFVLPLLCGVPVRNLCDGRFRAALDLLPSFLEWGTVTQCHPPFQTTRRHVIFDMRLSGSASQYITHLQRVDADLLGKGLLRHPELLLSRNGNPQRILCCLARLRSQARLQCARVLTVDSRRTFLDQLVLRGGSGVGLLDPIYCDRVGVFVGRLLHSTGLERRDLGLFTLSYIPSLYTLLRPEECLEDSVIQKLLREGCQ

#### >AspPolV-1-NL1-5a_P0

MFVITPEGFFLIDSSITPSFELLILLLRSLVAFCISALRTSSVPYDVFFFNFHRSFLFVLPLLCGVPVRNLCDGRFRAALDLLTAFLEWGTVTQCHPPFQTTRRHVIFDMRLSGSASQYITHLQRVDADLLGKGLLRHPELLLSRNGNPQRILCCLARLRSQARLQCARVLTVDSRRTFLDQLVLRGGSGVGLLDPIYCDRVGVFVGRLLHSTGLERRDLGLFTLSYIPSLYTLLRPEECLEDSVIQKLLREGCQ

#### >OV-5-NL1-5b_P0

MFVITPEGFFLIDSSITPSFELLILLLRSLVAFCFSAQRTSSVPDDVFFYNFRRSFLFVLPLLCGVPVRYFSDGRFRATLNLLPAFLEWGIVTQCHPPFQITKRHVIFDMRLFGSASRYITHLQRVDADLLGEGLLRHSELLLSRNGNPQRTLCCLAYLRAQARFQSARVLSVDSCRAFLDQLVLRGGSSVGLLDPVHCNRIGVLIGRLLHSTGLERRDLGLFTLSFIPSLYTLLRPEECLEDSVIQTLFREGCE

#### >AspPolV-1-NL1-9_P0

MFVITSEGFFLIDSSITPSFELLILLLRSLVAFCISALRTSSVPYDVFFFNFHRSFLFVLPLLCGVPVRNLCDGRFRAALDLLPSFLEWGTVTQCHPPFQTTRRHVIFDMRLSGSASQYITHLQRVDADLLGKGLLRHPELLLSRNGNPQRILCCLARLRSQARLQCARVLTVDSRRTFLYQLVLRGGSGVGLLDPIYCDRVGVFVGRLLHSTGLERHDLGLFTLSYVPSLYTLLRPEECLEDSVIQKLLREGCQ

#### >OV-5-NL1-24a_P0

MFVITPEGIFLIDSSISPSVELLTLLLRSLIAFCFSAQRTSSVPDDVFFYNFRRSFLFVLPLLCGVPVRYFSDGRFRATLNLLPAFLEWGIVTQCHPPFQITKRHVIFDMRLFGSASRYITHLQRVDADLLGEGLLRHSELLLSRNGNPQRTLCCLAYLRAQARFQSARVLSVDSCRAFLDQLVLRGGSSVGLLDPVHCNRIGVLIGRLLHSTGLERRDLGLFTLSFIPSLYTLLRPEECLEDSVIQTLFREGCE

#### >OV-5-NL1-24b_P0

MFVITPEGIFLIDSSISPSVELLTLLLRSLIAFCFSAQRTSSVPDDVFFYNFRRSFLFVLPLLCGVPVRYFSDGRFRATLNLLPAFLEWGIVTQCHPPFQITKRHVIFDMRLFGSASRYITHLQRVDADLLGEGLLRHSELLLSRNGNPQRTLCCLAYLRAQARFQSARVLSVDSCRAFLDQLVLRGGSSVGLLDPVHCNRIGVLIGRLLHSTGLERRDLGLFTLSFIPSLYTLLRPEECLEDSVIQTLFREGCE

#### >OV-5_P0

MFVITPEGIFLIDSSISPSVELLVLLLRSLIAFCFSAQRTSSVPDDVFFYNFRRSFLFVLPLLCGVPVRYFSDGRFRATLDLLPALLGWGIVTQCHPPFQITKRHVIFDMRLSGSASRYITHLQRVDADLLGEGLLRYPELLLSRNGNPQRKLCCLARLRARARFQSTRVLSVDSCRTFLDQLVLRGGGSVGLLDPVHCNRIGVLIGRLLHSTGLERRDLGLFTLSFIPSLYTLLRPEECLEDSIIQTLLREGCE

#### >PolV-M_P0

MFVITPEGIFLIDSSISPSVELLVLLLRSLIAFCFSAQRTSSVPDDVFFYNFRRSFLFVLPLLCGVPVRYFSDGRFRATLDLLPALLEWGIVTQCHPPFQTTKRHVIFDMRLSGSASRYITHLQRVDADLLGEGLLRYPELLLSRNGNPQRKLCCLARLRARARFQSTRVLSVDSCRTFLDQLVLRGGGSVGLLDPVHCNRIGVLIGRLLHSTGLERRDLGLFTLSFIPSLYTLLRPEECLEDSIIQTLLREGCE

#### >AspPolV-1-NL1-14_P1

MSSSSTFIVLFFLFSLSSVVSPSATSATGDFAPLSTFSPLFWNGALLPNATLLSRLPDAMSFSTCACPDPPLNTSLTYNELMRIFWEKVSSDTQSFSSRAMATLSEYSVALLASARKHASNALESLLWIPVALFWTSSYYAGVVVWVCLTQYTVIALAYLSVGCFIALAWNAAIWVCSRFPIFLLCTPFYVLRSVWKILSSKSFSVKVVNEKAVDGYVDYSIPQDPPRGSTVRLLYPNGNPLGYATCVRLFNGENALITSYHCVVEEGALVHSTRTGNKLPLSLFKPLYEDQRGDLSIMCGPPNWEGLMGCKGVHAVTCDRLGRGPATFFVYDNSWKARSAQITGRYEDFAQVLSNTEPGVSGAGYFNGKTLLGVHKGHTDVAEHNFNLMSPLPCLPGLTAPIYVYETTNVQGRIFDEDFVIQFKKGSWFDQMIEHAENHDVLPPLVRRPDGSLRFADEKSLPIEFSGNGAGSAVCPPNEPVKTPSAPPPMPSPSLLGSFVAPSSDTPVTVGVSSNALQEGVLKALVAKFDFQKVEQNVAELIAAKMLSRPQRTRGTRGRKRSNSKTTSPPSTSGQSQPPPKPQASKPSAGSPRTTTPPQNGKANGGKSSPGNIPSWVRKPQASAGRGLAQKRS

#### >AspPolV-1-NL1-2_P1

MSSSSTFIVLFFLFSLSSVVSPSATSATGDFAPLSTFSPLFWNGALLPNATLLSRLPDAMSFSTCACPDPPLNTSLTYNELMRIFWEKVSSDTQSFSSRAMATLSEHSVALLASARKHASNALESLLWIPVALFWTSLYYAGVVVWVCLTQYTVIALAYLSVGCFIALAWNAVIWVCSRFPIFLLCTPFYVLRS

VWKILSSKSFSVKVVNEKAVDGYVDYSIPQDPPRGSTVRLLYPNGNPLGYATCVRLFNGENALITSYHCVVEEGALVHSTRTGNKLPLSLFKPLYEDQRGDLSIMCGPPNWEGLMGCKGVHAVTCDRLGRGPATFFVYDNSWKARSAQIAGRYEDFAQVLSNTEPGVSGAGYFNGKTLLGVHKGHTDVAEHNFNLMSPLPCLPGLTAPIYVYETTNVQGRIFDEDFVIQFKKGSWFDQMIEHAENHDVLPPLVRRPDGSLRFADEKSLPIEFSGNGAGSAVCPPNEPVKTPSAPPPMPSPSLLGSFVAPSSDTPVTVGVSSNALQEGVLKALVAKFDFQKVEQNVAELIAAKMLSRPQRTRGTRGRKRSNSKTTSPPSTSGQSQPPPKPQASKPSAGSPRTTTPPQNGKANGGKSSPGNIPSWVRKPQASAGRGLAQRRS

#### >AspPolV-1-NL1-3_P1

MSSSSTFIVLFFLFSLSSVVSPSATSATGDFAPLSTFSPLFWNGALLPNATLLSRLPDAMSFSTCACPDPPLNTSLTYNELMRIFWEKVSSDTQSFSSRAMATLSEYSVALLASARKHAFNALESLLWIPVALFWTSLYYAGVVVWVCLTQYTVIALAYLSVGCFIALAWNAVIWVCSRFPIFLLCTPFYVLRSVWKILSSKSFSVKVVNEKAVDGYVDYSIPQDPPRGSTVRLLYPNGNPLGYATCVRLFNGENALITSYHCVVEEGALVHSTRTGNKLPLSLFKPLYEDQRGDLSIMCGPPNWEGLMGCKGVHAVTCDRLGRGPATFFVYDNSWKARSAQIAGRYEDFAQVLSNTEPGVSGAGYFNGKTLLGVHKGHTDVAEHNFNLMSPLPCLPGLTAPIYVYETTNVQGRIFDEDFVIQFKKGSWFDQMIEHAENHDVLPPLVRRPDGSLRFADEKSLPIEFSGNGAGSAVCPPNEPVKTPSAPPPMPSPSLLGSFVAPSSDTPVTVGVSSNALQEGVLKALVAKFDFQKVEQNVAELIAAKMLSRPQRTRGTRGRKRSNSKTTSPPSTSGQSQPPPKPQASKPSAGSPRTTTPPQNGKANGGKSSPGNIPSWVRKPQASAGRGLAQKRS

#### >AspPolV-1-NL1-5a_P1

MSSSSTFIVLFFLFSLSSVVSPSATSATGDSAPLSTFSPLFWNGALLPNATLLSRLPDAMSFSTCACPDPPLNTSLTYNELMRIFWEKVSFDTQSFSSRATATLSEYSVALLASARKHASNALESLLWIPVALFWTSLYYAGVVVWVCLTQYTVIALAYLSVGCFIALAWNAAIWVCSRFPIFLLCTPFYVLRSVWKILSSKSFSVKVVNEKAVDGYVDYSIPQDPPRGSTVRLLYPNGNPLGYATCVRLFNGENALITSYHCVVEEGALVHSTRTGNKLPLSLFKPLYEDQRGDLSIMCGPPNWEGLMGCKGVHAVTCDRLGRGPATFFVYDNSWKARSAQIAGRYEDFAQVLSNTEPGVSGAGYFNGKTLLGVHKGHTDVAEHNFNLMSPLPCLPGLTAPIYVYETTNVQGRIFDEDFVIQFKKGSWFDQMIEHAENHDVLPPLVRRPDGSLRFADEKSLPIEFSGNGAGSAVCPPNEPAETPSAPPPMPTPSLLGSFVAPPSDTPATVGVSSNALQEGVLKALVAKFDFQKVEQNVAELIAAKMLSRPQRTRGTRGRRRSNSKTTSPPSTSGQSQPPLKPQASKPLAGSPRTTTPPQNERASGGKSSPGNIPSWVRKPKASAGQGPAQKRS

#### >OV-5-NL1-5b_P1

MSSSTISVVLFFLFSLFFVASPSATSATGASAPLSTFSPLFWNGALLPNATLLSRLPNDMSSSTCVCSDPPLVISLTYNELMRIFSEKVCSDTRSFSSRAMATLSEHSVALLTSAHKHASRALESFLWIPVALFWTSLYYVGVVVWVCLTQYTAIALAYLLVGCFIALVWNAAIWVCSRFPLFLLCTPFYVLKSVWKTLSFKRSSVKVVNEKAVDGYVDYSIPQDPPRGSTVRLLYPNGNPLGYATCVRLFNGENALITSYHCVIEEGVLVHSTRTGNKLPLSLFKPLYEDQRGDLSIMCGPPNWEGLMGCKGVHAVTCDRLGRGPATFFVYDNSWKARSAQIAGRYEDFAQVLSNTEPGVSGAGYFNGKTLLGVHKGHTDVAEHNFNLMSPLPCLPGLTAPIYVYETTNVQGRIFDEDFVVQFKKGSWFDQMIEHAENHDVLPPLVRRPDGSLRFADEKSLPAKFSGNEAGSAVCPPNEPVKTPSAPPPMPSPSLLGSFVAPPSDTP

#### >AspPolV-1-NL1-9_P1

MSSSSTFIVLFFLFSLSSVVSPSATSATGDFAPLSTFSPLFWNGALLPNATLLSRLPDAMSFSTCACPDPPLNTSLTYNELMRIFWEKVSSDTQSFSSRAMATLSEYSVALLASARKHASNALESLLWIPVALFCTSLYYAGVVVWVCLTQYTVIALAYLSAGCFIALAWNAMIWVCSRFPMFLLCTPFYVLRSVWKILSSKSFSVKVVNEKAVDGYVDYSIPQDPPRGSTVRLLYPNGNPLGYATCVRLFNGENALITSYHCVVEEGALVHSTRTGNKLPLSLFKPLYEDQRGDLSIMCGPPNWEGLMGCKGVHAVTCDRLGRGPATFFVYDNSWKARSAQIAGRYEDFAQVLSNTEPGVSGAGYFNGKTLLGVHKGHTDVAEHNFNLMSPLPCLPGLTAPIYVYETTNVQGRIFDEDFVIQFKKGSWFDQMIEHAENHDVLPPLVRRPDGSLRFADEKSLPIEFSGNGAGSAVCPPNEPVKTPSAPPPMPSPSLLGSFVAPSSDTPVTVGVSSNALQEGVLKALVAKFDFQKVEQNVAELIAAKMLSRPQRTRGTRGRKRSNSKTTSPPSTSGQSQPPPKPQASKPSAGSPRTTTPPQNGKANGGKSSPGNIPSWVRKPQASAGRGLAQKRS

#### >OV-5-NL1-24a_P1

MSSSTISVVLFFLFSLFFVASPSATSATGASAPLSTFSPLFWNGALLPNATLLSRLPNDMSSSTCVCSDPPLVISLTYNELMRIFSEKVCSDTRSFSSRAMATLSEHSVALLTSAHKHASRALESFLWIPVALFWTSLYYVGVVVWVCLTQYTAIALAYLLVGCFIALVWNAAIWVCSRFPLFLLCTPFYVLKSVWKTLSFKRSSVKVVNEKAVDGYVDYSIPQDPPRGSTVRLLYPNGNPLGYATCVRLFNGENALITSYHCVIEEGVLVHSTRTGNKLPLSLFKPLYEDQRGDLSIMCGPPNWEGLMGCKGVHAVTCDRLGRGPATFFVYDNSWKARSAQIAGRYEDFAQVLSNTEPGVSGAGYFNGKTLLGVHKGHTDVAEHNFNLMSPLPCLPGLTAPIYVYETTNVQGRIFDEDFVVQFKKGSWFDQMIEHAENHDVLPPLVRRPDGSLRFADEKSLPAKFSGNEAGSAVCPPNEPVKTPSAPPPMPSPSLLGSFVAPPSDTPATVGVSSNALQEGVLKALVAKFDFQKVEQNVAELIAAKMLSRPQRTRGTRGRRRSNSKTTSPPSTSGQSQPPLKPQASKPLAGSPRTTTPPQNERASGGKSSPGNIPSWVRKPKASAGQGPAQKRS

#### >OV-5-NL1-24b_P1

MSSSTISVVLFFLFSLFFVASPSATSATGASAPLSTFSPLFWNGALLPNATLLSRLPNDMSSSTCVCSDPPLVISLTYNELMRIFSEKVCSDTRSFSSRAMATLSEHSVALLTSAHKHASRALESFLWIPVALFWTSLYYVGVVVWVCLTQYTAIALAYLLVGCFIALVWNAAIWVCSRFPLFLLCTPFYVLKSVWKTLSFKRSSVKVVNEKAVDGYVDYSIPQDPPRGSTVRLLYPNGNPLGYATCVRLFNGENALITSYHCVIEEGVLVHSTRTGNKLPLSLFKPLYEDQRGDLSIMCGPPNWEGLMGCKGVHAVTCDRLGRGPATFFVYDNSWKARSAQIAGRYEDFAQVLSNTEPGVSGAGYFNGKTLLGVHKGHTDVAEHNFNLMSPLPCLPGLTAPIYVYETTNVQGRIFDEDFVTQFKKGSWFDQMIEHAENHDVLPPLVRRPDGSLRFADEKSLPIEFSGNGAGSAVCPPNEPAETPSAPPPMPSPSLLGSFVAPSSDTPVTVGVSSNALQEGVLKALVAKFDFQKVEQNVAELIAAKMLSRPQRMRGTRGRKRSNSKTTSPPSTSGQSQPPPKPQASKPSAGSPRTTTPPQNGKANGGKSSPGNIPSWVRKPQASAGRGLAQKRS

#### >OV-5_P1

MSSSTISVVLFFLFSLFFVASPSATSATGASAPLSTFSPLFWDGALLPNATLLSRLPNDMSSSTCVCPDPPLGTSLTYNELMRIFSEKACSDIRSFSSRAMATLSENFVALLASAHEHASRALESFLWIPVALFWTSLYYVGVAVWVCLTQYTAIALAYLSVGCFIALAWNAAIWVCSRFPLFLLCTPFYVLRSVWKTLSFKRSSVKVVNEKAVDGYVDYSIPQDPPRGSTVRLLYPNGNPLGYATCVRLFNGENALITSYHCVIEEGVLVHSTRTGNKLPLSLFKPLYEDQRGDLSIMCGPPNWEGLMGCKGVHAVTCDRLGRGPATFFVYDNSWKARSAQIAGRYEDFAQVLSNTEPGVSGAGYFSGKTLLGVHKGHTDVAEHNFNLMSPLPCLPGLTAPIYVYETTNVQGRIFDEDFVIQFKKGSWFDKMIEHAENHDELPPLVRRPDGSLRFADEKSPSPKFSGNEAGSAVCPPNEPVKTPSAPPPMPSPSLLGSFVAPPSDTPATVGVSSNALQEGVLKALVAKFDFQKVEQNVAELIAAKMLSRPQRTRGTRGRKRSNSKTTSPPSTSGQSQPPLKPQASKPLAGSPRTTTPPQNERASGGKSSPGNIPSWVRKPKASAGQGPAQKRS

#### >PolV-M _P1

MSSSTISVVLFFLFSLFFVASPSATSATGASAPLSTFSPLFWNGALLPNATLLSRLPNDMSSSTCVCPDPPLGTSLTYNELMRIFSEKACSDIRSFSSRAMATLSENFVALLASAHEHASRALESFLWIPVALFWTSLYYVGVAVWVCLTQYTAIALAYLSVGCFIALAWNAAIWVCSRFPLFLLCTPFYVLRSVWKTLSFKRSSVKVVNEKAVDGYVDYSIPQDPPRGSTVRLLYPNGNPLGYATCVRLFNGENALITSYHCVIEEGVLVHSTRTGNKLPLSLFKPLYEDQRGDLSIMCGPPNWEGLMGCKGVHAVTCDRLGRGPATFFVYDNSWKARSAQIAGRYEDFAQVLSNTEPGVSGAGYFNGKTLLGVHKGHTDVAEHNFNLMSPLPCLPGLTAPIYVYETTNVQGRIFDEDFVIQFKKGSWFDKMIEHAENHDELPPLVRRPDGSLRFADEKSPSPKFSGNEAGSAVCPPNEPVKTPSAPPPMPSPSLLGSFVAPPSDTPATVGVSSNALQEGVLKALVAKFDFQKVEQNVAELIAAKMLSRPQRTRGTRGRKRSNSKTTSPPSTSGQSQPPLKPQASKPLAGSPRTTTPPQNERASGGKSSPGNIPSWVRKPKASAGQGPAQKRS

#### >AspPolV-1-NL1-14_P2

EIPPNRVLGKRSRQRRLPTKRACQDPVSPPPDALPLPSRKLCRSVLRHPCDRRRFFERLAGRSLKGVGSEVRLPEGRAECRRINRCENALPSSANAWYPREEAKQFQNYFASLYKWSESAPSEAPGFKTVGRLPSYYHPSPKRESKWGKKLTRQYPELGEKTTGFGWPGAGAEAELKSLQLQAARWLQRAETAIIPNQADRERVIARTVDAYCAVHTEVPSCARTNELTWPTFLEDFKRAICSLEVDAGIGIPYIAYGLPTHRGWVENPSLLPVLAQLTFARLQKLANVSFEALSAEELVRCGLCDPIRLFIKMEPHKQSKLDEGRYRLIMSVSLVDQLVARVLFQNQNQREIALWRVIPSKPGFGLSTDEQVADFLACLAKVVGTTPTEACSCWRELMIPTDCSGFDWSVAAWMLEDEMEVRNRLTINNTELCRRLRAGWTHCIANSVLCLSDGTLLAQTVPGVQKSGSYNTSSSNSRIRVMAAFHAGASWAMAMGDDALESTDSQIPVYKNLGFKVEVAEQLEFCSHVFVRPDLAIPTGTNKMLYKLLFGYDPECGNLEVLSNYIAACWSILNELRSDPELVSRLYSWLIPTQSQKNTAE

#### >AspPolV-1-NL1-2_P2

EIPPNRVLGKRSRQRRLPTKRACQDPVSPPPDALPLPSRKLCRSVLRHPCDRRRFFERLAGRSLKGVGSEVRLPEGRAECRRINRCENALPSSANAWYPREEAKQFQNYFASLYKWSESAPSEAPGFKTVGRLPSYYHPSPKRESKWGKKLTRQYPELGEKTTGFGWPGAGAEAELKSLQLQAARWLQRAETAIIPNQADRERVIARTVDAYCAVHTEVPSCARTNELTWPTFLEDFKRAICSLEVDAGIGIPYIAYGLPTHRGWVENPSLLPVLAQLTFARLQKLANVSFEALSAEELVRCGLCDPIRLFIKMEPHKQSKLDEGRYRLIMSVSLVDQLVARVLFQNQNQREIALWRAIPSKPGFGLSTDEQVADFLACLAKVVGTTPTEACSCWRELMIPTDCSGFDWSVAAWMLEDEMEVRNRLTINNTELCRRLRAGWTHCIANSVLCLSDGTLLAQTVPGVQKSGSYNTSSSNSRIRVMAAFHAGASWAMAMGDDALESTDSQIPVYKNLGFKVEVAEQLEFCSHVFVRPDLAIPTGTNKMLYKLLFGYDPECGNLEVLSNYIAACWSILNELRSDPELVSRLYSWLIPTQSQKNTAE

#### >AspPolV-1-NL1-3_P2

EIPPNRVLGKRSRQRRLPTKRACQDPVSPPPDALPLPSRKLCRSVLRHPCDRRRFFERLAGRSLKGVGSEVRLPEGRAECRRINRCENALPSSANAWYPREEAKQFQNYFASLYKWSESAPSEAPGFKTVGRLPSYYHPSPKRESKWGKKLTRQYPELGEKTTGFGWPGAGAEAELKSLQLQAARWLQRAETAIIPNQADRERVIARTVDAYCAVHTEVPSCARTNELTWPTFLEDFKRAICSLEVDAGIGIPYIAYGLPTHRGWVENPSLLPVLAQLTFARLQKLANVSFEALSAEELVRCGLCDPIRLFIKMEPHKQSKLDEGRYRLIMSVSLVDQLVARVLFQNQNQREIALWRVIPSKPGFGLSTDEQVADFLACLAKVVGTTPTEACSCWRELMIPTDCSGFDWSVAAWMLEDEMEVRNRLTINNTELCRRLRAGWTHCIANSVLCLSDGTLLAQTVPGVQKSGSYNTSSSNSRIRVMAAFHAGASWAMAMGDDALESTDSQIPVYKNLGFKVEVAEQLEFCSHVFVRPDLAIPTGTNKMLYKLLFGYDPECGNLEVLSNYIAACWSILNELRSDPELVSRLYSWLIPTQSQKNTAE

#### >AspPolV-1-NL1-5a_P2

EIPPNRVLGKRSRQRRLPTKRACRDPVSPPPDAHPLPSRKLCRSTLRHPCDRRRFFKRLAGRCPQGVGSEVRLPEGRAECRRIDCCENALPSSANAWYPREETKQFQNYFASLYKWSEPAPSEAPGFKTVGRLPSYYHPSPKRESKWGKKLTRQHPELGEKTQGFGWPGAGAEAELKSLQLQAARWLQRAETAIIPNQADRERVIARTVDAYCAVHTEVPSCARTNELTWPTFLEDFKRAICSLEVDAGIGIPYIAYGLPTHRGWVENPSLLPVLAQLTFARLQKLANVSFEALSAEELVRCGLCDPIRLFIKMEPHKQSKLDEGRYRLIMSVSLVDQLVARVLFQNQNQREIALWRAIPSKPGFGLSTDEQVADFLACLAKVVGTTPTEACSRWRELMIPTDCSGFDWSVAAWMLEDEMEVRNRLTINNTELCRRLRAGWTHCIANSVLCLSDGTLLAQTVPGVQKSGSYNTSSSNSRIRVMAAFHAGASWAMAMGDDALESTDSQIPVYKNLGFKVEVAEQLEFCSHIFVRPDLAIPTGTNKMLYKLLFGYDPECGNLEVLSNYIAACWSILNELRSDPELVSRLYSWLIPTQSQKNTAE

#### >AspPolV-1-NL1-9_P2

EIPPNRVLGKRSRQRRLPTKRACQDPVSPPPDALPLPSRKLCRSVLRHPCDRRRFFERLAGRSLKGVGSEVRLPEGRAECRRINRCENALPSSANAWYPREEAKQFQNYFASLYKWSESAPSEAPGFKTVGRLPSYYHPSPKRESKWGKKLTRQYPELGEKTTGFGWPGAGAEAELKSLQLQAARWLQRAETAIIPNQADRERVIARTVDAYCAVHTEVPSCARTNELTWPTFLEDFKRAICSLEVDAGIGIPYIAYGLPTHRGWVENPSLLPVLAQLTFARLQKLANVSFEALSAEELVRCGLCDPIRLFIKMEPHKQSKLDEGRYRLIMSVSLVDQLVARVLFQNQNQREIALWRAIPSKPGFGLSTDEQVADFLACLAKVVGTTPTEACSCWRELMIPTDCSGFDWSVAAWMLEDEMEVRNRLTINNTELCRRLRAGWTHCIANSVLCLSDGTLLAQTVPGVQKSGSYNTSSSNSRIRVMAAFHAGASWAMAMGDDALESTDSQIPVYKNLGFKVEVAEQLEFCSHVFVRPDLAIPTGTNKMLYKLLFGYDPECGNLEVLSNYIAACWSILNELRSDPELVSRLYSWLIPTQSQKNTAE

#### >OV-5-NL1-24a_P2

KIPSRQILGKRSWQRRLPTKRACQDPVSPPPDALPLPSRKLCRSTLRHPCDRRRFFKRLAGRCPQGVGSEVRLPEGRAECRRIDCCENALPSSANAWYPREETKQFQNYFASLYKWSEPAPSEAPGFKTVGRLPSYYHPSPKRESKWGKKLTRQHPELGEKTQGFGWPGAGAEAELKSLQLQAARWLQRAETATIPNQADRERVIARTVDAYCAVHTEVPSCARTNELTWPTFLEDFKRAICSLEVDAGIGIPYIAYGLPTHRGWVENPSLLPVLAQLTFARLQKLANVSFEALSAEELVRCGLCDPIRLFIKMEPHKQSKLDEGRYRLIMSVSLVDQLVARVLFQNQNQREIALWRAIPSKPGFGLSTDEQVADFLACLAKVVGTTPTEACSRWRELMIPTDCSGFDWSVAAWMLEDEMEVRNRLTINNTELCRRLRAGWTHCIANSVLCLSDGTLLAQTVPGVQKSGSYNTSSSNSRIRVMAAFHAGASWAMAMGDDALESTDSQIPVYKNLGFKVEVAEQLEFCSHIFVRPDLAIPTGTNKMLYKLLFGYDPECGNLEVLSNYIAACWSILNELRSDPELVSRLYSWLIPTQSQKNTAE

#### >OV-5-NL1-24b_P2

KIPSRQILGKRSWQRRLPTKRACQDPVSPPPDALPLPSRKLCRSTLRHPCDRRRFFKRLAGRCPQGVGSEVRLPEGRAECRRIDCCENALPSSANAW YPREETKQFQNYFASLYKWSEPAPSEAPGFKTVGRLPSYYHPSPKRESKWGKKLTRQHPELGEKTQGFGWPGAGAEAELKSLQLQAARWLQRAETAT IPNQADRERVIARTVDAYCAVHTEVPSCARTNELTWPTFLEDFKRAICSLEVDAGIGIPYIAYGLPTHRGWVENPSLLPVLAQLTFARLQKLANVSFEALSAEELVRCGLCDPIRLFIKMEPHKQSKLDEGRYRLIMSVSLVDQLVARVLFQNQNQREIALWRAIPSKPGFGLSTDEQVADFLACLAKVVGTTPTEACSRWRELMIPTDCSGFDWSVAAWMLEDEMEVRNRLTINNTELCRRLRAGWTHCIANSVLCLSDGTLLAQTVPGVQKSGSYNTSSSNSRIRVMAAFHAGASWAMAMGDDALESTDSQIPVYKNLGFKVEVAEQLEFCSHIFVRPDLAIPTGTNKMLYKLLFGYDPECGNLEVLSNYIAACWSILNELRSDPELVSRLYSWLIPTQSQKNTAE

#### >OV-5_P2

KIPSRQILGKRSWQRRLPTKRACQDPVSPPPDALPLPSRKLCRSTLRHPCDRRRFFKRLAGRCPQGVGSEVRLPEGRAECRRIDCCENALPSSANAW YPREETKQFQNYFASLYKWSEPAPSEAPGFKTVGRLPSYYHPSPKRESKWGKKLTRQHPELGEKTQGFGWPGAGAEAELKSLQLQAARWLQRAETAT IPNQADRERVIARTVDAYCAVHTEVPSCARTNELTWPTFLEDFKRAICSLEVDAGIGIPYIAYGLPTHRGWVENPSLLPVLAQLTFARLQKLANVSF EALSAEELVRCGLCDPIRLFIKMEPHKQSKLDEGRYRLIMSVSLVDQLVARVLFQNQNQREIALWRAIPSKPGFGLSTDEQVADFLACLAKVVGTTPTEACSRWRELMIPTDCSGFDWSVAAWMLEDEMEVRNRLTINNTELCRRLRAGWTHCIANSVLCLSDGTLLAQTVPGVQKSGSYNTSSSNSRIRVMAA FHAGASWAMAMGDDALESTDSQIPVYKNLGFKVEVAEQLEFCSHIFVRPDLAIPTGTNKMLYKLLFGYDPECGNLEVLSNYIAACWSILNELRSDPE LVSRLYSWLIPTQSQKNTAE

#### >PolV-M_P2

EVPFPEVLGKRSRQRRLPTKRACQDPVSPPPDALPLPSRKLCRSTLRHPCDRRRFFKRLAGRCPQGVGSEVRLPEGRAECRRIDCCENALPSSANAWYPREEAKQFQNYFASLYKWSEPAPSEAPGFKTVGRLPSYYHPSPKRESKWGKKLTRQHPELGEKTQGFGWPGAGAEAELKSLQLQAARWLQRAETAT

IPNQADRERVIARTVDAYCAVHTEVPSCARANELTWPTFLEDFKRAICSLEVDAGIGIPYIAYGLPTHRGWVENPSLLPVLAQLTFARLQKLANVSFEALSAEELVRCGLCDPIRLFIKMEPHKQSKLDEGRYRLIMSVSLVDQLVARVLFQNQNQREIALWRAIPSKPGFGLSTDEQVADFLACLAKVVGTTPTEACSRWRELMIPTDCSGFDWSVAAWMLEDEMEVRNRLTINNTELCRRLRAGWTHCIANSVLCLSDGTLLAQTVPGVQKSGSYNTSSSNSRIRVMAAFHAGASWAMAMGDDALESTDSKIPVYKDLGFKVEVAEQLEFCSHIFVRPDLAIPTGTNKMLYKLLFGYDPECGNLEVFSNYIAACWSILNELRSDPELVSRLYSWLIPTQSQKNTAE

#### >AspPolV-1-NL1-14_P3

MNTGGARSRNGRRRVRLPRRLQRRRPAQPIVVVSGPNQIRRRRRRRGNSSRRGANAAPGRGGSRETFVFTKDDLAGNTSGSLTFGPSLSECPAFATG ILKAYHEYRISQCTLEFISEAPSTASGSIAYELDAHCKISSLSSKINKFGITKGGKRSFAASKINGIEWHDSSEDQFRLLYKGNGASNIAGSFRVTI VVATQNPK

#### >AspPolV-1-NL1-2_P3

MNTGGARSRNGRRRVRLPRRLQRRRPAQPIVVVSGPNQIRRRRRRRGNSSRRGANAAPGRGGSRETFVFTKDDLAGNTSGSLTFGPSLSECPAFATG ILKAYHEYRISQCTLEFISEAPSTASGSIAYELDAHCKISSLSSKINKFGITKGGKRSFAASKINGIEWHDSSEDQFRLLYKGNGASNIAGSFRVTI VVATQNPK

#### >AspPolV-1-NL1-3_P3

MNTGGARSRNGRRRVRLPRRLQRRRPAQPIVVVSGPNQIRRRRRRRGNSSRRGANAAPGRGGSRETFVFTKDDLAGNTSGSLTFGPSLSECPAFATG ILKAYHEYRISQCTLEFISEAPSTASGSIAYELDAHCKISSLSSKINKFGITKGGKRSFAASKINGIEWHDSSEDQFRLLYKGNGASNIAGSFRVTI VVATQNPK

#### >AspPolV-1-NL1-5a_P3

MNTGGARSKNGRRRVRLPRRLQRRRPAQPIVVVSGPNQIRRRRRRRGNSSRRGANAASGRGGSRETFVFTKDDLAGNTSGSLTFGPSLSECPAFATG ILKAYHEYRISQCTLEFISEAPSTASGSIAYELDAHCKISSLSSKINKFGITKGGKRSFAASKINGIEWHDSSEDQFRLLYKGNGASNIAGSFRVTI VVATQNPK

#### >AspPolV-1-NL1-9_P3

MNTGGARSRNGRRRVRLPRRLQRRRPAQPIVVVSGPNQIRRRRRRRGNSSRRGANAAPGRGGSRETFVFTKDDLAGNTSGSLTFGPSLSECPAFATG ILKAYHEYRISQCTLEFISEAPSTASGSIAYELDAHCKISSLSSKINKFGITKGGKRSFAASKINGIEWHDSSEDQFRLLYKGNGASNIAGSFRVTI VVATQNPK

#### >OV-5-NL1-24a_P3

MNTGGARSRNGRRRVRLPRRLQRRRPAQPIVVVSGPNQIRRRRRRRGNSSRRGANAASGRGGSRETFVFTKDDLAGNTSGSLTFGPSLSECPAFATG ILKAYHEYRISQCTLEFISEAPSTASGSIAYELDAHCKISSLSSKINKFGITKGGKRSFAASKINGIEWHDSSEDQFRLLYKGNGASNIAGSFRVTI VVATQNPK

#### >OV-5-NL1-24b_P3

MNTGGARSRNGRRRVRLPRRLQRRRPAQPIVVVSGPNQIRRRRRRRGNSSRRGANAASGRGGSRETFVFTKDDLAGNTSGSLTFGPSLSECPAFATG ILKAYHEYRISQCTLEFISEAPSTASGSIAYELDAHCKISSLSSKINKFGITKGGKRSFAASKINGIEWHDSSEDQFRLLYKGNGASNIAGSFRVTI VVATQNPK

#### >OV-5_P3

MNTGGARSRNGRRRVRLPRRLQRRRPAQPIVVVSGPNQIRRRRRRRGNSSRRGANAAPGRGGSRETFVFTKDDLAGNTSGSLTFGPSLSECPAFATG ILKAYHEYRISQCTLEFVSEAPSTASGSIAYELDAHCKISSLSSKINKFGITKGGKRSFAASKINGIEWHDSSEDQFRLLYKGNGASNIAGSFRVTI VVATQNPK

#### >PolV-M_P3

MNTGGARSRNGRRRVRLPRRLQRRRPAQPIVVVSGPNQIRRRRRRRGNSSRRGANAAPGRGGSRETFVFTKDDLAGNTSGSLTFGPSLSECPAFATG ILKAYHEYRISQCTLEFVSEAPSTASGSIAYELDAHCKISSLSSKINKFGITKGGKRSFAASKINGIEWHDSSEDQFRLLYKGNGASNIAGSFRVTI VVATQNPK

#### >AspPolV-1-NL1-14_P3-5

MNTGGARSRNGRRRVRLPRRLQRRRPAQPIVVVSGPNQIRRRRRRRGNSSRRGANAAPGRGGSRETFVFTKDDLAGNTSGSLTFGPSLSECPAFATG ILKAYHEYRISQCTLEFISEAPSTASGSIAYELDAHCKISSLSSKINKFGITKGGKRSFAASKINGIEWHDSSEDQFRLLYKGNGASNIAGSFRVTI VVATQNPK*VDGEPGPSPGPDPPPPAPAPQPHKHERFIAYVGIPMLTIQAKENDDGIILRSLGPQRMKYIEDENQNYTNIDSQYYSQTSVSAVPMYY FNVPSGKWSVDISCEGYQCTSSTTDPNRGRSDGLIAYSNSSDDYWNVGEADGVIITKLTNDNTYRMGHPDLEVNSCHFRDGQLLERDATISFHVEASSDGRFFLIGPAIQKMEKYNYTISYGEWTDRDMELGLITVILDEHLEDSGSRFGRIQRRKTRDGHVHLTSIAERQPLTQTPRSLRVLDTGSRHKSLSPERARSPSPLREPSPPPPENPEMWQVDWHAPKSGEPFAPLPQDFKRPESALKGILREARGNTGNYERDYADKENWDPSTLEDPTRFGTNWTSSQIAEAQEQTDAKRKTRFSLVPRSFKGGSSLGGGSLTGGSLRNTLRGRLEKLTAEQLIQYHRIKQQQGSTVAQMFLTDCLGDG

#### >AspPolV-1-NL1-2_P3-5

MNTGGARSRNGRRRVRLPRRLQRRRPAQPIVVVSGPNQIRRRRRRRGNSSRRGANAAPGRGGSRETFVFTKDDLAGNTSGSLTFGPSLSECPAFATG ILKAYHEYRISQCTLEFISEAPSTASGSIAYELDAHCKISSLSSKINKFGITKGGKRSFAASKINGIEWHDSSEDQFRLLYKGNGASNIAGSFRVTI VVATQNPK*VDGEPGPSPGPDPPPPAPAPQPHKHERFIAYVGIPMLTIQAKENDDGIILRSLGPQRMKYIEDENQNYTNIDSQYYSQTSVSAVPMYY FNVPSGKWSVDISCEGYQCTSSTTDPNRGRSDGLIAYSNSFDDYWNVGEADGVIITKLTNDNTYRMGHPDLEVNSCHFRDGQLLERDATISFHVEASSDGRFFLIGPAIQKMEKYNYTISYGEWTDRDMELGLITVILDEHLEDSGSRFGRIQRRKTRDGHVHLTSIAERQPLTQTPRSLRVLDTGSRHKSLSPERARSPSPLREPSPPPPENPEMWQVDWHAPKSGEPFAPLPQDFKRPESALKGVLRESRGNTGNYERDYADKENWDPSTLEDPTRFGTNWTSSQIAEAQEQTDAKRKTRFSLVPRSFKGGSSLGGGSLTGGSLRNTLRGRLEKLTAEQLIQYHRIKQQQGSTVAQMFLTDCLGDG

#### >AspPolV-1-NL1-3_P3-5

MNTGGARSRNGRRRVRLPRRLQRRRPAQPIVVVSGPNQIRRRRRRRGNSSRRGANAAPGRGGSRETFVFTKDDLAGNTSGSLTFGPSLSECPAFATG ILKAYHEYRISQCTLEFISEAPSTASGSIAYELDAHCKISSLSSKINKFGITKGGKRSFAASKINGIEWHDSSEDQFRLLYKGNGASNIAGSFRVTI VVATQNPK*VDGEPGPSPGPDPPPPAPAPQPHKHERFIAYVGIPMLTIQAKENDDGIILRSLGPQRMKYIEDENQNYTNIDSQYYSQTSVSAVPMYY FNVPSGKWSVDISCEGYQCTSSTTDPNRGRSDGLIAYSNSSDDYWNVGEADGVIITKLTNDNTYRMGHPDLEVNSCHFRDGQLLERDATISFHVEASSDGRFFLIGPAIQKMEKYNYTISYGEWTDRDMELGLITVILDEHLEDSGSRFGRIQRRKTRDGHVHLTSIAERQPLTQTPRSLRVLDTGSRHKSLSPERARSPSPLREPSPPSPENPEMWQVDWHAPKSGEPFAPLPQDFKRPESALKGVLREARGNTGNYERDYADKENWDPSTLEDPTRFGTNWTSSQIAEAQEQTDAKRKTRFSLVPRSFKGGSSLGGGSLTGGSLRNTLRGRLEKLTAEQLIQYHRIKQQQGSTVAQMFLTDCLGDG

#### >AspPolV-1-NL1-5a_P3-5

MNTGGARSKNGRRRVRLPRRLQRRRPAQPIVVVSGPNQIRRRRRRRGNSSRRGANAASGRGGSRETFVFTKDDLAGNTSGSLTFGPSLSECPAFATG ILKAYHEYRISQCTLEFISEAPSTASGSIAYELDAHCKISSLSSKINKFGITKGGKRSFAASKINGIEWHDSSEDQFRLLYKGNGASNIAGSFRVTI VVATQNPK*VDGEPGPSPGPDPPPPAPAPQPRKHERFIAYVGIPMLTIQAKKNDDGIILRSLGPQRMKYIEDENQNYTNIDSQYYSQTNVSAVPMYY FNVPAGKWSVDVSCEGYQCTSSTTDPNRGRSDGLIAYSNNSDDYWNVGEADGVIITKLTNDNTYRMGHPDLEVNSCHFRDGQLLERDATISFHVEASSDGRFFLIGPAIQKMEKYNYTISYGEWTDRDMELGLITVILDEHLEDSGSRFGKIQRRKTRDGHVRLTSVAERQPLVQTPRPLRVLDTGSRRKSLSPERANSPPPLREPSPPPPEDPELWQVGWYTPKSGEPLAPLPQDFKRPESVLKGILRESRGNTGDYEKDYADKENWDPSTLEDPTRFGTNWTSSQIAEAQEQTDAKRKTRFSLVPRSFKGGSSISGGSLSGGSLRSTLRSRLEKLTAEQQIQYHRIKQQQGATTAQMFLSDCLGDG

#### >AspPolV-1-NL1-9_P3-5

MNTGGARSRNGRRRVRLPRRLQRRRPAQPIVVVSGPNQIRRRRRRRGNSSRRGANAAPGRGGSRETFVFTKDDLAGNTSGSLTFGPSLSECPAFATG ILKAYHEYRISQCTLEFISEAPSTASGSIAYELDAHCKISSLSSKINKFGITKGGKRSFAASKINGIEWHDSSEDQFRLLYKGNGASNIAGSFRVTI VVATQNPK*VDGEPGPSPGPDPPPPVPAPQPHKHERFIAYVGIPMLTIQAKENDDGIILRSLGPQRMKYIEDENQNYTNIDSQYYSQTSVSAVPMYY FNVPSGKWSVDISCEGYQCTSSTTDPNRGRSDGLIAYSNSSDDYWNVGEADGVTITKLTNDNTCRMGHPDLEVNSCHFRDGQLLERDATISFHVEASSDGRFFLIGPAIQKMEKYNYTISYGEWTDRDMELGLITVILDEHLEDSGSRFGRIQRRKTRDGHVHLTSIAERQPLTQTPRSLRVLDTGSRHKSLSPERARSPSPLREPSPPPPENPEIWQVDWHTPKSGEPFAPLPQDFKRPESALKGVLREARGNTGNYERDYADKENWDPSTLEDPTRFGTNWTSSQIAEAQEQTDAKRKTRFSLVPRSFKGGSSLGGGSLTGGSLRNTLRGRLEKLTAEQLIQYHRIKQQQGSSVAQMFLTDCLGDG

#### >OV-5-NL1-24a_P3-5

MNTGGARSRNGRRRVRLPRRLQRRRPAQPIVVVSGPNQIRRRRRRRGNSSRRGANAASGRGGSRETFVFTKDDLAGNTSGSLTFGPSLSECPAFATG ILKAYHEYRISQCTLEFISEAPSTASGSIAYELDAHCKISSLSSKINKFGITKGGKRSFAASKINGIEWHDSSEDQFRLLYKGNGASNIAGSFRVTI VVATQNPK*VDGEPGPSPGPDPPPPAPAPQPRKHERFIAYVGIPMLTIQAKENDDGIILRSLGPQRMKYIEDENQNYTNIDSQYYSQTNVSAVPMYY FNVPAGKWSVDVSCEGYQCTSSTTDPNRGRSDGLIAYSDNSDDYWNVGEADGVIITKLTNDNTYRMGHPDLEVNSCHFRDGQLLERDATISFHVEAS SDGRFFLIGPAIQKMEKYNYTISYGEWTDRDMELGLITVILDEHLEDSGSRFGKIQRRKTRDGHVRLTSVAERQPLVQTPRPLRVLDTGSRRKSLSP ERANSPPPL

#### >OV-5-NL1-24b_P3-5

MNTGGARSRNGRRRVRLPRRLQRRRPAQPIVVVSGPNQIRRRRRRRGNSSRRGANAASGRGGSRETFVFTKDDLAGNTSGSLTFGPSLSECPAFATG ILKAYHEYRISQCTLEFISEAPSTASGSIAYELDAHCKISSLSSKINKFGITKGGKRSFAASKINGIEWHDSSEDQFRLLYKGNGASNIAGSFRVTI VVATQNPK*VDGEPGPSPGPDPPPPAPAPQPRKHERFIAYVGIPMLTIQAKENDDGIILRSLGPQRMKYIEDENQNYTNIDSQYYSQTNVSAVPMYY FNVPAGKWSVDVSCEGYQCTSSTTDPNRGRSDGLIAYSDNSDDYWNVGEADGVIITKLTNDNTYRMGHPDLEVNSCHFRDGQLLERDATISFHVEASSDGRFFLIGPAIQKMEKYNYTISYGEWTDRDMELGLITVILDEHLEDSGSRFGKIQRRKTRDGHVRLTSVAERQPLVQTPPPLRVLDTGSRRKSLSPERANSPPPRREPSPPPPEDPELWQGGWYTPKSGEPLAPLPQDFKRPESVLKGIQRESRGNTGNYEKDYADKENRDPSTLEDPTRFGTNWTSSQIAEAQEQTDAKRKTRFSLVPRSFKGGSSISGGSLSGGSLRSTLRSRLEKLTAEQQIQYHRIKQQQGATTAQMFLSDCLGDG

#### >OV-5_P3-5

MNTGGARSRNGRRRVRLPRRLQRRRPAQPIVVVSGPNQIRRRRRRRGNSSRRGANAAPGRGGSRETFVFTKDDLAGNTSGSLTFGPSLSECPAFATG ILKAYHEYRISQCTLEFVSEAPSTASGSIAYELDAHCKISSLSSKINKFGITKGGKRSFAASKINGIEWHDSSEDQFRLLYKGNGASNIAGSFRVTI VVATQNPK*VDGEPGPSPGPDPPPPAPAPQPHKHERFIAYVGIPMLTIQAKENDDGIILRSLGQQRMKYIEDENQNYTNIDSQYYSQTNVSATPMYY FNVPAGKWSVDISCEGYQSTSSTTDPNRGRSDGLIAYSNSSDDYWNVGEADGVVITKLTNDNTYRMGHPDLEVNSCHFRDGQLLERDATISFHVEASNNGRFFLIGPAIQKTAKYNYTISYGEWTDRDMELGLITVILDEHLEDSGSRFGRIQRRKTRDGHVHLTSVAERQPLIQTPRSSRVLDSGSRRKSLSPERARSPSPLREPSPPPPEDPELWQVDWHTPKPGEPFATLPQDFRRPESVLKGVLREARGNTGNYERDYADKENWDPSTLEDPTRFGTNWTSSQIAEAQEQTDAKRKTRFSLVPRSFKGGSSLGGGSLTGGSLRNALRGRLEKLTAEQLIQYHKIKQQQGSSAAQMFLTDCLGDG

#### >PolV-M_P3-5

MNTGGARSRNGRRRVRLPRRLQRRRPAQPIVVVSGPNQIRRRRRRRGNSSRRGANAAPGRGGSRETFVFTKDDLAGNTSGSLTFGPSLSECPAFATG ILKAYHEYRISQCTLEFVSEAPSTASGSIAYELDAHCKISSLSSKINKFGITKGGKRSFAASKINGIEWHDSSEDQFRLLYKGNGASNIAGSFRVTI VVATQNPK*VDGEPGPSPGPDPPPPAPAPQPHKHERFIAYVGIPMLTIQAKENDDGIILRSLGQQRMKYIEDENQNYTNIDSQYYSQTNVSATPMYY FNVPAGKWSVDISCEGYQSTSSTTDPNRGRSDGLIAYSNSSDDYWNVGEADGVVITKLTNDNTYRMGHPDLEVNSCHFRDGQLLERDATISFHVEASNNGRFFLIGPAIQKTAKYNYTISYGEWTDRDMELGLITVILDEHLEDSGSRFGKIQRRKTRDGHVHLTSVAERQPLIQTPRSSRVLDSGSRRKSLSPERARSPSPLREPSPPPPEDPELWQVDWHTPKPGEPFATLPQDFRRPESVLKGVLREARGNTGNYERDYADKENWDPSTLEDPTRFGTNWTSSQIAEAQEQTDAKRKTRFSLVPRSFKGGSSLGGGSLTGGSLRNALRGRLEKLTAEQLIQYHKIKQQQGSSAAQMFLTDCLGDG

#### >AspPolV-1-NL1-14_P4

MDVVEFDYHAASSGVGQLSQSLWSRDPTRFDAVDDEEEILHVEEQMQLQEEEALVKHLFSRRMTSRATPLEVSPSGRVFQSVQHSRQEFSKPTMSIVSHSVLWSSSPKPLPRRPVQSLMSWMHTAKSPLSPQKSTSLESLKEGRGALRRPKSTG

#### >AspPolV-1-NL1-2_P4

MDVVEFDYHAASSGVGQLSQSLWSRDPTRFDAVDDEEEILHVEEQMQLQEEEALVKHLFSRRMTSRATPLEVSPSGRVFQSVQHSRQEFSKPTMSIVSHSVLWSSSPKPLPRRPVQSLMSWMHTAKSPLSPQKSTSLESLKEGRGALRRPKSTG

#### >AspPolV-1-NL1-3_P4

MDVVEFDYHAASSGVGQLSQSLWSRDPTRFDAVDDEEEILHVEEQMQLQEEEALVKHLFSRRMTSRATPLEVSPSGRVFQSVQHSRQEFSKPTMSIVSHSVLWSSSPKPLPRRPVQSLMSWMHTAKSPLSPRKSTSLESLKEGRGALRRPKSTG

#### >AspPolV-1-NL1-5a_P4

MDVVEFDYHAASSGVGQLSQSLWSQDPTRFDAVDDEEEILHVEEQMQLQEEEALVKHSFSRRMTSRATPLEVSPSGRVFQSVQHSRQEFSKPTMSIVSHSVLWSSSPKPLPRRPVQSLMSWMHIAKSPLSPRKSTSLESPKEGRGALRRPKSTG

#### >AspPolV-1-NL1-9_P4

MDVVEFDYHAASSGVGQLSQSLWSRDPTRFDAVDDEEEILHVEEQMQLQEEEALVKHLFSRRMTSRATPLEVSPSGRVFQSVQHSRQEFSKPTMSIVSHSVLWSSSPKPLPRRPVQSLMSWMHTAKSPPSPQKSTSLESLKEGRGALRRPKSTG

#### >OV-5-NL1-24a_P4

MDVVEFDYHAASSGVGQLSQSLWSQDPTRFDAVDDEEEILHVEEQMQLQEEEALVKHSFSRRTTSRATPLEVSPSGRVFQSVQHSRQEFSKPTMSIVSHSVLWSSSPKPLPRRPVQSLMSWMHIAKSPLSPRKSTSLESPKEGRGALRRPKSTG

#### >OV-5-NL1-24b_P4

MDVVEFDYHAASSGVGQLSQSLWSQDPTRFDAVDDEEEILHVEEQMQLQEEEALVKHSFSRRTTSRATPLEVSPSGRVFQSVQHSRQEFSKPTMSIVSHSVLWSSSPKPLPRRPVQSLMSWMHIAKSPLSPRKSTSLESPKEGRGALRRPKSTG

#### >OV-5_P4

MDVAEFDYHAASSGVGQLSQSLWSQDPTRFDAVDDEEEILHVEEQMQLQEEEALVKHSFSRRMTSRATPLEVSPSGRVFQSVQHSRQEFSKPTMSIVSHSVLWSSSPKPLPRRPVQSLMSWMHTAKSPPSPRKLTSSESPKEGRGALRRPKSTG

#### >PolV-M_P4

MDVAEFDYHAASSGVGQLSQSLWSQDPTRFDAVDDEEEILHVEEQMQLQEEEALVKHSFSRRMTSRATPLEVSPSGRVFQSVQHSRQEFSKPTMSIV SHSVLWSSSPKPLPRRPVQSLMSWMHTAKSPPSPRKLTSSESPKEGRGALRRPKSTG

